# Corticotropin-releasing hormone signaling from prefrontal cortex to lateral septum supports social novelty preference

**DOI:** 10.1101/2022.03.15.484224

**Authors:** Noelia Sofia de León Reyes, Paula Sierra Díaz, Ramon Nogueira, Antonia Ruiz-Pino, Yuki Nomura, Christopher de Solis, Jay Schulkin, Arun Asok, Félix Leroy

## Abstract

Social preference, the decision to interact with one member of the same species over another, is a key feature of optimizing social interactions. In rodents, social preference relies on both extrinsic factors, such as sex, strain and kinship, and intrinsic ones, such as the memory of previous encounters, which favors interactions with novel compared to familiar animals (social novelty preference). At present, it is unclear which neuronal circuits guide social preferences and whether such circuits promote social interactions with the preferred individuals or suppress interactions with the non-preferred ones. Although both the infra-limbic area of the pre-frontal cortex (ILA) and the lateral septum (LS) have been shown to support social novelty preference, the neuronal circuits and molecular mechanisms by which these brain regions interact to regulate social interactions are unknown. Here, we identify a population of inhibitory neurons in ILA that express the neuropeptide corticotropin releasing hormone (CRH) and project to the rostro-dorsal region of LS (rdLS). Release of CRH from ILA in rdLS during interactions with familiar mice disinhibits rdLS neurons, thereby suppressing interactions with familiar mice and contributing to social novelty preference. We further demonstrate how the maturation of CRH expression during the first two post-natal weeks enables the developmental shift from a preference for littermates in juveniles to a preference for novel mice in adults.

## Introduction

Social preference, the decision to interact with one conspecific over another, is a feature displayed by gregarious animals which is critical to navigate their social space^1,2^. Adult rodents prefer to interact with their kin^3,4^, individuals from specific strains^5^ and members of the opposite sex^6–9^. In addition to innate factors (e.g., kin, strain, and sex), social preference is also influenced by social memory^10^, social hierarchy^9,11,12^ and the affective state of the conspecific^13^. Thus, adult rodents display social novelty preference, choosing to interact with novel individuals over familiar ones^10^. For the last two decades, social novelty preference has been used as a proxy to assess social memory^14–16^ but the neuronal circuits mediating social novelty preference and whether such circuits promote social interactions with the preferred individuals or prevent interactions with the non-preferred ones are unknown.

Memory-based social preferences, such as social novelty preference, have a developmental window^17^ and can change during the life of altricial animals. Young mice prefer their mother to an unfamiliar dam for the first few weeks^18^. After weaning, mice display reversed social preference for an unfamiliar dam over the mother^18^. Similarly, rat pups display a preference for their familiar siblings during the first 2 postnatal weeks, after which preference shifts toward novel pups^3,4^. Although the mechanisms that regulate these shifts remains elusive, the lateral septum (LS), a brain region associated with the regulation of motivated behaviors including social interactions^19^, is necessary for kinship/familiarity preference in young rats^3^ as well as for social novelty preference in adult rodents^19–21^. Moreover, the ventral aspect of medial prefrontal cortex (mPFC), the infra-limbic area (ILA), is known for its involvement in decision-making and recent evidence shows it responds to social stimuli^22–24^ and is necessary for social novelty preference^25,26^. Although PFC is known to project to LS^27^, how these regions integrate social memory cues and communicate to regulate social interactions and display a preference for the novel or familiar individual is still poorly understood.

Corticotropin-releasing hormone (CRH)^28^, a 41 amino acid peptide, regulates diverse processes ranging from homeostatic neuroendocrine mechanisms to memory^29^, including social behaviors in non-stressful context^30,31^. In human, CRH has been involved in psychiatric disorders associated with social deficits such as depression^32,33^ or social phobia^34^. In rodents, knocking-out the gene encoding CRH in mouse forebrain inhibitory neurons decreases social interaction^35^ and central administration of CRH inhibits social interactions in rats^36–40^. Moreover, application of the CRH receptor 1 agonist stressin-1 elicited the same effect^41^ suggesting that CRH effects on social interactions are mediated by the CRH receptor 1 (CRHR1). CRH also plays an important role in social novelty preference and social recognition^31,42,43^. Social recognition in rats is facilitated by over-expressing CRH^43^ or increasing CRH tone by pharmacological targeting of CRH receptors or binding proteins^31^. Importantly, administration of a CRH-binding protein ligand inhibitor during social recognition did not affect social interaction during the initial exposure but impaired performance during recall^31^.

Despite the evidence implicating CRH and its receptor in social interactions and social novelty preference, the neural circuitry mediating its release and targeted by its action remains unknown. Given that CRH is expressed in ILA^44^ and CRHR1 is expressed in LS^45^, we hypothesized that CRH release from PFC to LS is involved in social novelty preference.

We demonstrate through a combination of electrophysiological, chemogenetic, optogenetic, calcium recording and gene silencing techniques that the release of CRH from ILA neurons (ILA^CRH^ neurons) during familiar interactions into the rostro-dorsal region of LS (rdLS) suppresses social interaction between familiar mice. This circuit therefore controls familiarization (decrease in interaction as a novel rodent becomes familiar) and contributes to the social novelty preference exhibited by adult mice. In addition, the increase in ILA^CRH^ neurons density during the second postnatal week is responsible for a shift in the social preference of young mice from familiar to novel conspecifics.

## Results

### ILA^CRH^ cells project to rdLS

We injected *CRH-Cre* mice in ILA with a Cre-dependent adeno-associated virus (AAV) expressing membranous GFP and synaptophysin tagged with mRuby in order to visualize axons and terminals respectively (Fig. 1a-b). We observed GFP^+^ fibers in the rostral-dorsal section of the lateral septum (rdLS, Fig. 1c-d). A closer examination confirmed the presence of mCherry^+^ axon terminals in this region. We did not observe fibers going to other brain regions. To confirm this projection, we injected the retrograde marker CtB-488 in rdLS of *CRH-Cre* mice crossed with a Cre-dependent TdTomato reporter line (*CRH-Cre;Ai9* mice) to visualize CRH^+^ cells (Fig. 1f-g). CtB travelled retrogradely to ILA neurons mostly located in layer 2/3 (Fig. 1h-k). Some CtB^+^ cells also expressed CRH-TdTomato (Fig. 1i-j). The cells were found mainly in ILA layer 2/3 (Fig. 1l). Overall, these experiments show that ILA^CRH^ cells project to rdLS.

**Figure 1:**
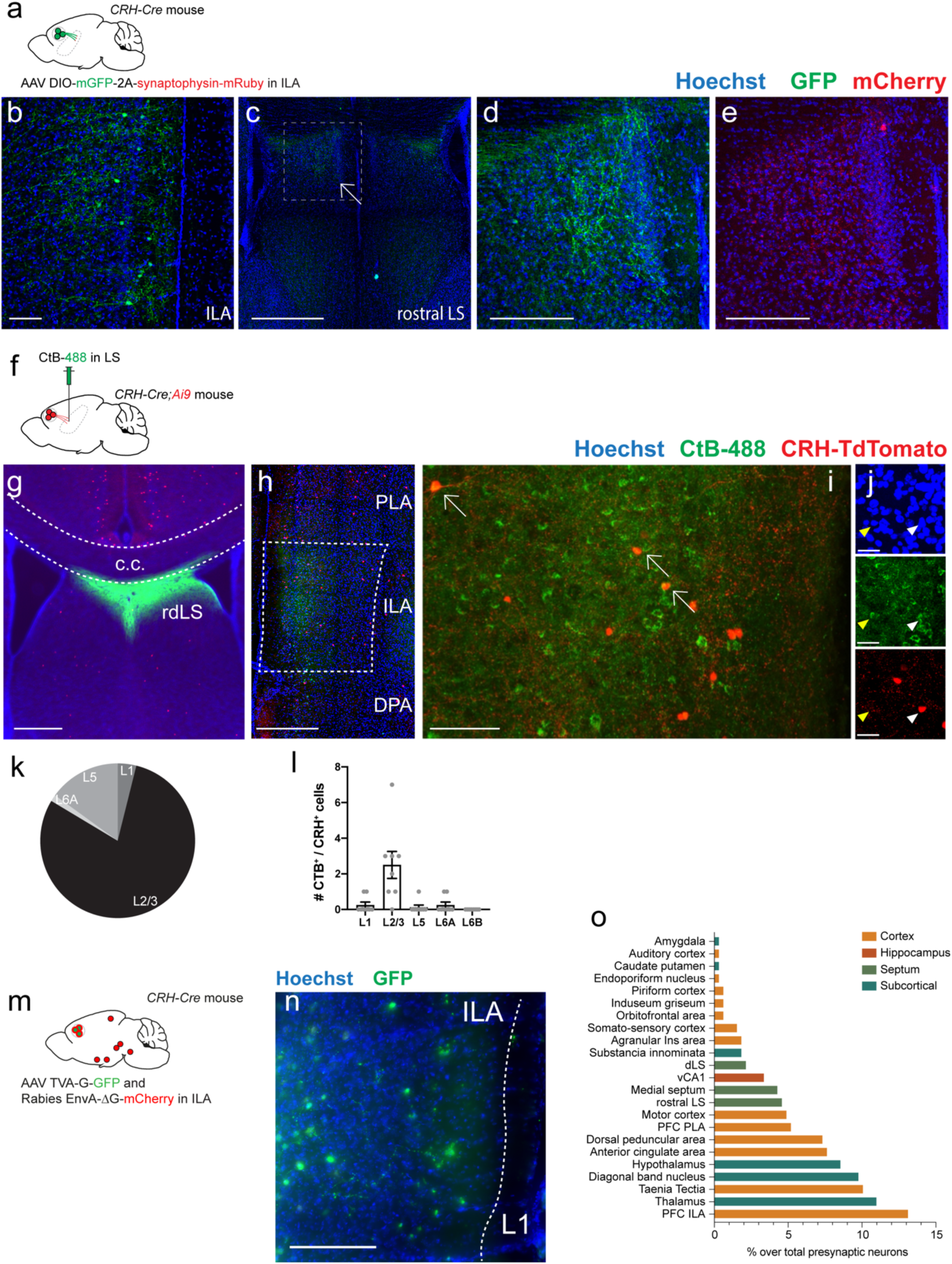
ILA^CRH^ cells project to the rostro-dorsal LS. **a.** *CRH-Cre* mice injected in ILA with AAV2/DJ hSyn.FLEX.mGFP.2A.Synatophysin-mRuby. **b-e.** Immunohistochemistry images of an ILA (b) and LS (c-e) section labelled for GFP (b-d) or mRuby (e). Scale bars: 100 µm (b), 500 µm (c) and 200 µm (d-e). **f.** *CRH-Cre;Ai9* mice injected in rdLS with CtB-488. **g.** Image of a coronal brain section containing the injection site. Scale bar: 400 µm. **h-j.** Images of coronal brain sections containing the mPFC. Scale bar: 400 µm (h), 200 µm (i) and 50 µm (j). **k.** Distribution of CtB^+^ cells in ILA. **l.** Number of CtB^+^/CRH^+^ cells per ILA layer. Each point is from a different section. N = 3 mice. Bar graph represent mean ± S.E.M. One-way ANOVA, *p* < 0.0001. **m.** *CRH-Cre* mice injected in ILA with AAV2/8 syn.DIO.TVA-2A-GFP-2A-B19G and rabies SAD-B19.EnvA.ΔG.mCherry. **n.** Immunohistochemistry images of an ILA section showing GFP-expressing “starter” cells. Scale bar: 200 µm. **o.** Relative distribution of presynaptic Rabies-infected mCherry^+^ cells in the brain. dLS: dorsal lateral septum. vCA1: ventral CA1 region. PLA: peri-limbic area of the PFC. DPA: dorsal peduncular area of the PFC. ACA: anterior cingulate area. ILA: infra-limbic area of the PFC. N = 3 mice.

We confirmed this result by injecting a Cre-dependent retrograde monosynaptic herpes simplex virus expressing GFP in rdLS of *CRH-Cre* mice (Extended Data Fig. 1). Consistent with our CtB-488 injections, 79% of CRH/GFP^+^ cells were located in the ILA with the rest being located in adjacent regions (Extended Data Fig. 1b-c). Within ILA, 66% of GFP^+^ cells were located in layer 2/3.

Using in situ hybridization markers for excitatory and inhibitory neurons, we found that 92% of ILA^CRH^ cells expressed the mRNA for glutamic acid decarboxylase 2 (*Gad2*) while only 3% expressed the mRNA for the vesicular glutamate transporter 1 (*Slc17a7*/VGlut1). Immunofluorescent labeling demonstrated that 89% of GFP^+^ ILA^CRH^ cells are positive for GABA, confirming the identity of these cells as GABAergic inhibitory neurons (Extended Data Fig. 1g-h). To dissect further the molecular identity of ILA^CRH^ cells, we labelled coronal brain sections from *CRH-Cre;Ai9* mice with PV and SST antibodies and found 3% and 8% of overlap respectively (Extended Data Fig. 1i-m). Overall, ILA^CRH^ cells projecting to rdLS are a sub-population of GABAergic cells.

Next, we asked which brain regions provide input to the population of ILA^CRH^ cells. We injected *CRH-Cre* mice in ILA with rabies helper AAVs and G-deleted pseudotyped rabies virus to label pre-synaptic neurons impinging on ILA^CRH^ cells (Fig. 1m-n). Rabies-mCherry^+^ cells were found throughout the brain. Preeminent inputs came from all mPFC regions, other cortical regions, hippocampal ventral CA1, dorsal thalamus, hypothalamus and medial and lateral septum (Fig. 1o & Extended Data Fig. 2).

### ILA^CRH^ cells regulate the duration of social interaction with a familiar mouse

Next, we used a chemogenetic approach in order to modulate the activity of ILA^CRH^ cells and probe their behavioral function. We injected *CRH-Cre* mice in ILA with Cre-dependent AAVs expressing an inhibitory DREADD tagged with mCherry (iDREADD) or mCherry only as a control (Fig. 2a-b). Three weeks later, mice were injected with the DREADD agonist CNO (5 mg/kg intra-peritoneally) 30 min prior to conducting the behavioral tests. Since a previous study^44^ associated CRH^+^ cells in the dorso-medial PFC area with anxiety, we first tested the mice in the open-arena to assess locomotion and anxiety (Extended Data Fig. 3a). Silencing ILA^CRH^ cells had no effect on the distance travelled, the time spent in the center or the surround or the ratio of time spent in the surround vs. center (Extended Data Fig. 2b-d). Next, we ran the mice for the elevated plus maze test of anxiety and found no effect on the number of entry or time spent in the open arms relative to the closed ones (Extended Data Fig. 2e-i). Finally, since glutamatergic cells in the PFC projecting to LS have been reported to be involved in food-seeking behavior, we also ran the mice for the anxiety-suppressed feeding behavior test, where a food-deprived mouse must venture into the center of an open-arena in order to eat (Extended Data Fig. 2j). Silencing ILA^CRH^ cells had no effect on the latency to feed, the duration or the number of entries into the food zone (Extended Data Fig. 2k-m).

**Figure 2:**
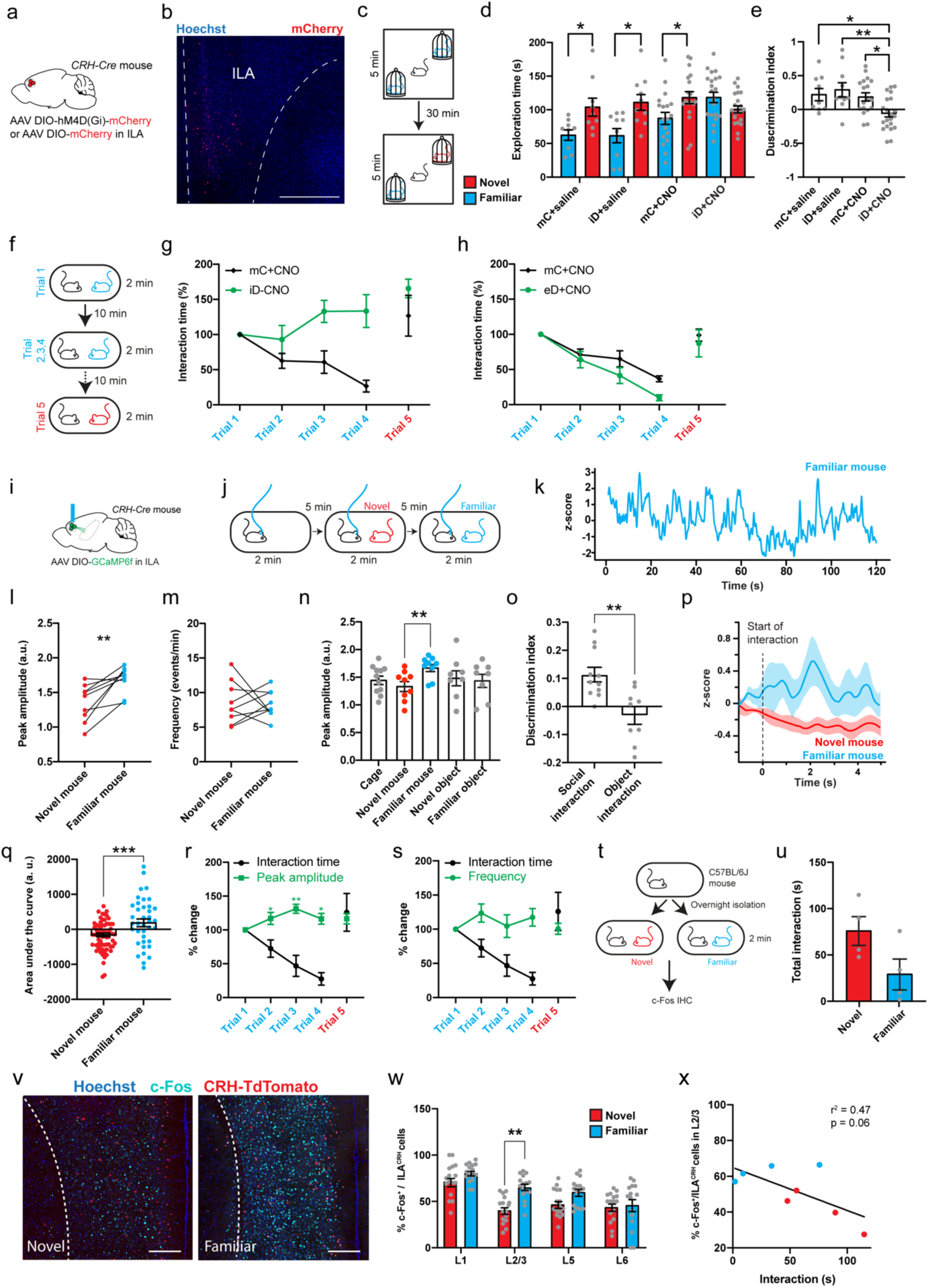
ILA^CRH^ cells are necessary for social memory and respond preferentially to familiar mice. **a.** *CRH-Cre* mice injected in ILA with AAV2/8 hSyn.DIO.hM4D(Gi)-mCherry (iDREADD) or AAV2/8 hSyn.DIO.mCherry. **b.** Immunohistochemistry image showing viral expression in the ILA. Scale bar: 500 µm. **c.** Schematic of the social novelty preference test. **d.** Interaction time with novel (red) or familiar (blue) mouse during trial 2 in mice expressing mCherry (mC) or hM4Di (iD). Both groups were injected with saline (left groups) or 5 mg/kg of the DREADD agonist CNO (right groups). Grey dots are different mice. Paired *t* tests: *p* = 0.04, *p* = 0.02, *p* = 0.03 and *p* = 0.2. **e.** Discrimination indexes for social novelty preference of the four groups during recall trial. One-sample *t* tests compared to 0: mC + saline, *p* = 0.04; iD + saline, *p* = 0.02; mC + CNO, *p* = 0.006; iD + CNO, *p* = 0.2. Two-way ANOVA: trial x injection F_1, 54_ = 4,795, *p* =0,03; virus F_1, 54_ = 7,068, *p* = 0,01; injection F_1, 54_ = 1,535, *p* = 0,2. Tukey’s multiple comparisons tests compared to the iD + CNO group: mC + saline, *p* = 0.04; iD + saline, *p* = 0.008; mC + CNO, *p* = 0.04. **f.** Schematic of the repetitive social presentation test. **g.** Normalized interaction times during social presentations (inhibitory DREADD-expressing mice and controls injected with CNO). 8 mice per group. Two-way ANOVA: trial x virus F_12,139_ = 2.09, *p* = 0.02; trial F_4,139_ = 17.21, *p* < 0.0001; virus F_3,139_ = 15.76, *p* < 0.0001. **h.** Normalized interaction times during repetitive social presentation test in *CRH-Cre* mice injected in ILA with AAV5 hSyn.DIO.hM3D(Gq)-mCherry (excitatory DREADD) or with AAV5 hSyn.DIO.mCherry as a control. 8 mice per group. Two-way ANOVA: trial x virus F_12,140_ = 0.96, *p* = 0.5; trial F_4,140_ = 34.21, *p* < 0.0001; virus F_3,140_ = 3.01, *p* = 0.03. **i.** *CRH-Cre* mice injected in ILA with AAV2/1 syn.FLEX.GCaMP6f and implanted with an optical ferrule over ILA. **j.** Schematic of the fiber-photometry recording experiment. **k.** Example trace of recording during presentation of a familiar mouse. **l.** Average peak amplitude of the z-score during social presentations. Each dot is a different recording session using 2 mice. Paired *t* test: *p* = 0.005. **m.** Frequency of calcium events during presentation of a novel or familiar mouse. Paired *t* test: *p* = 0.8. **n.** Average peak amplitudes during each type of presentation. Kruskal-Wallis test: F_4,36_ = 4.991, *p* = 0.07. Multiple comparisons tests: cage vs. familiar mouse, *p* = 0.3; novel vs. familiar mouse, *p* = 0.02; novel object vs. familiar mouse, *p* = 0.3; familiar object vs. familiar mouse, *p* = 0.3. **o.** Discrimination indexes for familiarity preference calculated from z-scores during mouse or object presentation. Each point is one recording session. N = 4 mice. One-sample *t* tests compared to 0: *p* = 0.001 and *p* = 0.3 respectively. Unpaired *t* test between groups: *p* = 0.002. **p.** Peri-stimulus time histogram during social interaction with novel or familiar mouse. **q.** Area under the curve during familiar and novel mouse interaction. Each point is an interaction. N = 4 mice. Unpaired *t* test between groups: *p* = 0.0003. **r.** Fiber-photometry recording during repetitive social presentation test (10 sessions in 5 mice). One-sample *t* test compared to: *p* = 0.06 (trial 2), *p* =. 0.002 (trial 3) and *p* = 0.04 (trial 4). **s.** Frequency of calcium events during repetitive social presentation test (10 sessions in 5 mice). **t.** *CRH-Cre;Ai9* mice were presented with novel or familiar mice after overnight isolation before being processed for immunohistochemistry. **u.** Interaction times following 2 min social presentation. **v.** Immunohistochemistry images of c-Fos labelling in ILA. Scale bars: 200 µm. **w.** Percentage of ILA^CRH^ cells positive for c-Fos per layer. Each point corresponds to each side of 2 sections. 4 mice per group. Unpaired *t* test, *p* < 0.0001. **x.** Percentage of ILA^CRH^ cells positive for c-Fos in layer 2/3 vs. interaction time during social interaction with novel (red) or familiar (blue) mouse. Each point represents one mouse. For the entire figure, bar graphs represent mean ± S.E.M.

The above results argue against a prominent function of ILA^CRH^ cells in locomotion, anxiety or feeding behaviors. We next tested whether these cells could regulate social interactions. The mPFC is known to regulate sociability, social preference, social hierarchy as well as emotion discrimination^13,25,26^ but it remains unclear whether specific sub-regions or populations control different facets of social interactions. First, we silenced ILA^CRH^ cells and assessed the mice’s sociability (preference for a novel mouse compared to a novel object, Extended Data Fig. 4a)^14^. Both groups exhibited a strong preference for the mouse compared to the object (Extended Data Fig. 4b-c).

Next, we tested whether ILA^CRH^ cells regulate social novelty preference (Fig. 2c). A subject mouse was exposed to two novel stimulus mice inside wire cup cages in opposite corners of a squared open arena. After 5 min exploring both mice (learning trial), the subject mouse was removed from the arena, placed into an empty housing cage, and one of the two stimulus mice was replaced by a third (novel) mouse. After a 30 min inter-trial interval the subject mouse was reintroduced in the arena (recall trial). Social novelty preference is manifest when the subject mouse spends more time exploring the novel stimulus mouse compared to the familiar one during the recall trial. We also quantified the preference for exploring the novel mouse by calculating a discrimination index (DI), representing the percentage of extra time the subject mouse spent with the novel compared to the familiar (see methods).

We measured the social novelty preference behavior of mice in which iDREADD-expressing ILA^CRH^ cells were silenced from the start of the task by injecting CNO systemically 30 min prior to the learning trial. To rule out off-target effects of CNO or of iDREADD expression alone, we examined three control groups of mice: 1. mice injected with CNO expressing mCherry in ILA^CRH^ cells, 2. Mice injected with saline expressing iDREADD in ILA^CRH^ cells; 3. Mice injected with saline expressing mCherry in ILA^CRH^ cells. During recall, the three control groups (mCherry + saline, iDREADD + saline, mCherry + CNO) exhibited a higher interaction time with the novel mouse compared to the familiar one (Fig. 2d), which translated into a high discrimination index preference for the novel mouse (Fig. 2e), indicating intact social novelty preference. However, in the test group in which ILA^CRH^ cells were silenced, the subject mice explored the novel and familiar mice to the same extent (Fig. 2d). As a result, the discrimination index for social novelty preference was not different from zero (Fig. 2e). During learning or recall, the total exploration time of the mice was similar across groups (Extended Data Fig. 4d-e), suggesting that ILA^CRH^ cell silencing does not affect the motivation to explore. Overall, this experiment shows that ILA^CRH^ cells are necessary for social novelty preference.

How ILA^CRH^ cells regulates social novelty preference is however unclear. Are they important for social memory or rather for downstream processes such as social novelty preference that utilize social memory cues? Because PFC is involved in executive functions^26^, we supposed that ILA^CRH^ cells leverage these social memory cues to guide social preference by regulating social interactions. Do ILA^CRH^ cells support social novelty preference by promoting interactions with the novel mouse or by suppressing interactions with the familiar one? Because familiar and novel mice are presented simultaneously during the recall trial, this test is not adapted to answer this point. Some insights can however be gleaned from measuring the social interactions with 2 novel mice during the learning phase of test. Mice in the test group explored novel conspecifics to the same extent than control groups (Extended Data Fig. 4f), suggesting that silencing ILA^CRH^ cells does not impair social interaction with novel animals. We therefore tested the role of ILA^CRH^ cells during repetitive social interactions with a single novel mouse as it becomes familiar.

We used the repetitive social presentation test (also known as the habituation/dishabituation test), where a sex- and age-matched novel mouse is presented 4 times to the test mouse (Fig. 2f). Control mice showed a progressive decrease in interaction time with repeated presentations of the mouse (Fig. 2g). The interaction time jumped back to its initial level, when a novel mouse was presented in the final 5^th^ trial, demonstrating that the decreased interaction was not due to fatigue or loss of engagement in the task. In contrast, mice expressing iDREADD showed no decrease in interaction during the repeated presentations, suggesting that ILA^CRH^ cells are necessary for social familiarization (Fig. 2g). We repeated the experiments injecting saline instead of CNO and both groups exhibited a steady decrease in interaction (Extended Data Fig. 4g). Next, we asked whether over-activating ILA^CRH^ cells could promote social familiarization by performing the repetitive social presentation test using mice expressing an excitatory DREADD in ILA. Increasing the activity of ILA^CRH^ cells with CNO facilitated the decrease in social interaction (Fig. 2h), which was not observed when mice were injected with saline (Extended Data Fig. 4h), indicating that ILA^CRH^ cells can bidirectionally modulate the interaction time with familiar mice. Taken together, these experiments suggest that ILA^CRH^ cells repress social interaction with a familiar mouse and are necessary for social familiarization.

To confirm DREADD modulated ILA^CRH^ cells activity, *CRH-Cre* mice expressing excitatory or inhibitory DREADD in ILA were injected with CNO or saline before presenting them with a familiar animal (Extended Data Fig. 5b). We measured the overlap between c-Fos and mCherry expression in ILA (Extended Data Fig. 5a). iDREADD-expressing mice given CNO exhibited less c-Fos/mCherry^+^ cells compared to saline control showing ILA^CRH^ cells silencing (Extended Data Fig. 5c). On the contrary, eDREADD-expressing mice injected with CNO showed an increased overlap showing ILA^CRH^ cells activation.

Do ILA^CRH^ cells specifically control social interactions or does it extend to objects as well? We performed tests of object recognition memory and repetitive object presentation while silencing the cells and found no effect (Extended Data Fig. 6), indicating that ILA^CRH^ are specifically involved in regulating social interactions.

If ILA^CRH^ cells regulate interaction with a familiar mouse, we can expect the cells to be more active when the mouse interacts with familiar mice than with novel ones. To test this prediction, we performed fiber-photometry of ILA^CRH^ cells. We injected the ILA of *CRH-Cre* mice with a Cre-dependent AAV expressing the calcium sensor GCaMP6f and implanted an optical ferrule above ILA (Fig. 2i & Extended Data Fig. 7a). Subject mice were presented with familiar and novel mice in random order and the activity of the cells recorded (Fig. 2j). We automatically detected the peaks of calcium transients and measured their average amplitude and frequency during each trial (Fig. 2k). The peak amplitudes were higher during familiar compared to novel mouse presentation, but we saw no difference in the frequency of events (Fig. 2l-m). Overall, these observations suggest that a neuronal population of ILA^CRH^ cells increase their activity during familiar social interactions. The order of presentation did not affect this result (Extended Data Fig. 7b). To determine whether ILA^CRH^ cell activity differs during novel and familiar mouse presentation, we also trained linear classifiers to discriminate between interactions with a novel or familiar mouse using our fiberphotometry recordings. We implemented 2 classifiers using either individual recording sessions (Extended Data Fig. 7c, left) or a meta-session pooling all sessions (pseudo-simultaneous, Extended Data Fig. 7c, right)^46^. For each classifier, we also computed chance levels using permutation tests (grey areas). Most individual recording sessions yielded a decoding performance above chance with an average 68% accuracy. The pseudo-simultaneous data yielded a decoding performance even higher (79%). This shows that the population of ILA^CRH^ cells can code for social familiarity.

We also presented novel and familiar objects and saw no change in activity compared to baseline (Fig. 2n) or between novel and familiar object (Extended Data Fig. 7d-f). Using the peak amplitudes, we calculated the discrimination indexes (DI) for familiarity preference following object or social presentation (Fig. 2o). DI of social interaction showed a strong preference for social familiarity, unlike the one for object interaction. We also calculated the peri-stimulus time histogram using the start of social interaction to synchronize episodes and found that familiar mouse presentation elicited a large increase of the response while presenting a novel mouse did not (Fig. 2p-q). We then recorded ILA^CRH^ cells during the repetitive social presentation test and found the peak amplitude to increase during familiarization (Fig. 2r). The frequency of events however remained stable (Fig. 2s), similar to what was observed previously (Fig. 2m).

As a further assay of neural activity, we measured the expression of the immediate-early gene c-Fos. *CRH-Cre;Ai9* mice were presented with a novel or familiar mouse for 2 min (Fig. 2t). As expected, mice interacted more with novel than familiar mice (Fig. 2u). Mice were perfused 1 hour later and processed for c-Fos immunohistochemistry in order to count the number of ILA^CRH^ and rdLS neurons expressing c-Fos (Fig. 2v and 4k respectively). Despite shorter interactions, ILA^CRH^ cells exhibited higher c-Fos expression following encounters with familiar compared to novel mice (Fig. 2w), similar to what was reported for the entire ILA cell population^22^. This increase was significant only in ILA^CRH^ cells located in the layer 2/3, which are the ones projecting to rdLS (Fig. 1). The activation of ILA^CRH^ cells negatively correlated with the amount of social interaction (Fig. 2x). Overall, our experiments demonstrate that ILA^CRH^ cells are more active during interaction with a familiar mouse than with a novel one.

### CRH release from ILA in rdLS regulates the duration of social interaction with a familiar mouse

How could ILA^CRH^ cells suppress social interactions with a familiar mouse and regulate social novelty preference? ILA^CRH^ projects to rdLS and LS regulates social interactions^19^, which led us to ask whether CRH release from ILA in rdLS is necessary for familiarization. We designed 3 different shRNAs against *Crh* and had their efficacy tested in cell cultures overexpressing CRH. We selected the most efficient one and recombined it into a Cre-dependent plasmid which also expressed mCherry (see methods). Plasmids were packaged into AAVs and injected into the ILA of *CRH-Cre* mice (Fig. 3a, top). To assess the efficacy of the shRNA approach, we performed in situ hybridization to measure CRH and mCherry mRNA levels in brain slices with the ILA (Fig. 3a, bottom). CRH^+^ cells expressing *mCherry* (viral marker) showed a 4-fold decrease in the intensity of *Crh* labelling compared to nearby non-infected CRH^+^ cells that did not express *mCherry* (Fig. 3a-b). No changes were seen in mice expressing the scrambled shRNA (Fig. 3a-b). These results indicate that our strategy to reduce *Crh* level in ILA^CRH^ neurons is both specific and efficient.

**Figure 3:**
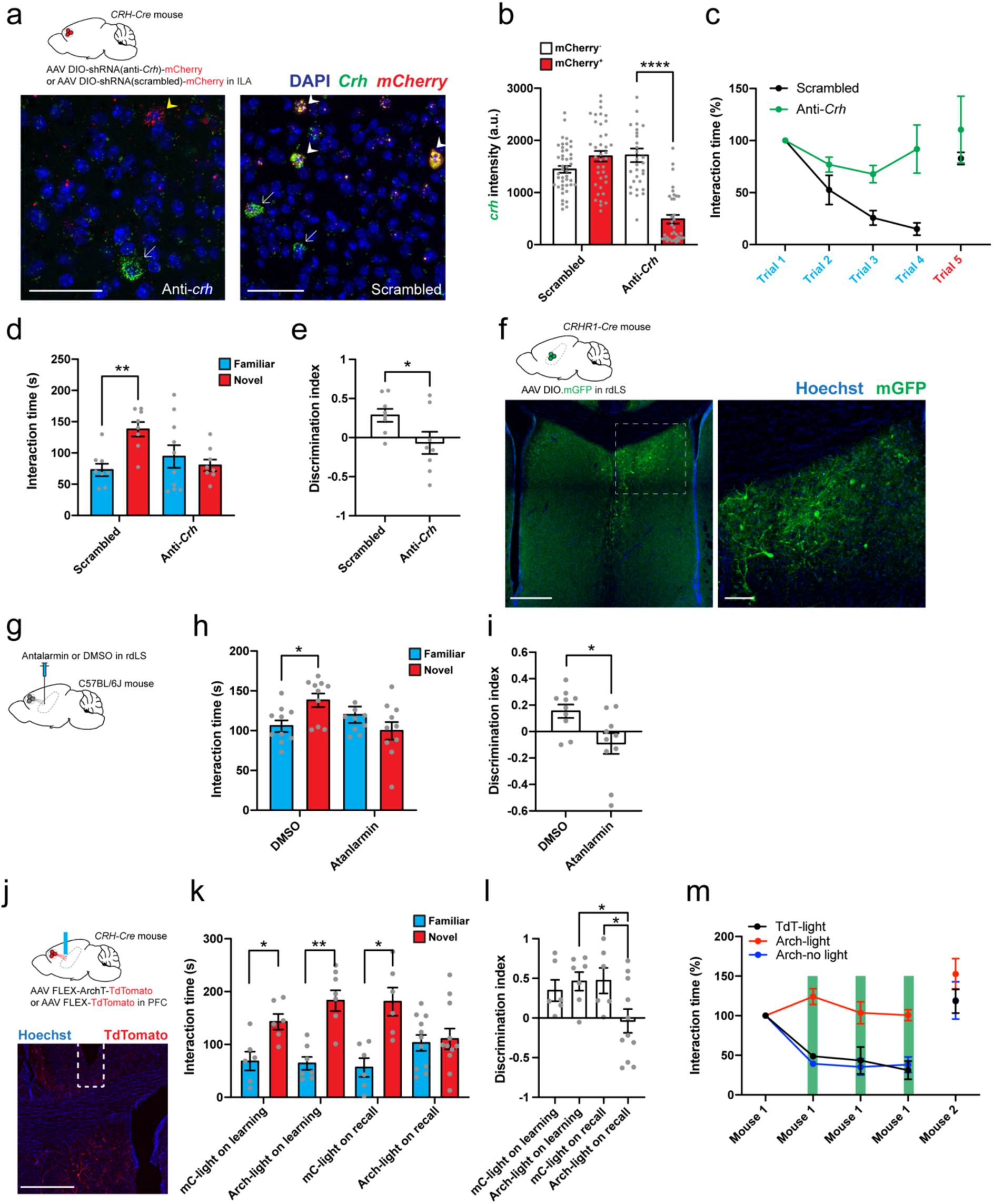
CRH release from ILA to rdLS regulates social memory. **a.** *CRH-Cre* mice injected in ILA with AAV2/9 CMV-DIO-(mCherry-U6)-shRNA(anti-*Crh*) to downregulate *Crh* or control AAV2/9 CMV-DIO-(mCherry-U6)-shRNA(scrambled) (top). In situ hybridization images of ILA slices expressing shRNA against *crh* (left) or a scrambled shRNA (right) labelled for *mCherry* and *Crh*. Yellow arrowhead denotes a Cre-, mCherry- and anti-*Crh* shRNA-expressing neuron with reduced levels of *Crh*. White arrowheads denote *Crh*^+^ neurons that express the scrambled shRNA and *Crh.* White arrows denote neurons that do not express the virus (*mCherry*^-^) with intact levels of *Crh*. Scale bars: 50 µm. **b.** Quantification of *Crh* expression in cells using in situ hybridization images in ILA slices from mice injected with AAV expressing scrambled or anti-*Crh* shRNAs. In each slice neurons were classified as to whether they were uninfected or infected with virus based on *mCherry* expression (3 mice per group; each point is from a different neuron). Note reduction in *Crh* expression in neurons infected with anti-*Crh* shRNA. 2-way ANOVA; shRNA x viral expression F_1, 144_ = 61.34, *p* < 0.0001; shRNA F_1, 144_ = 24.80, *p* < 0.0001; viral expression F_1, 144_ = 26.75, *p* < 0.0001 followed by Tukey’s post-hoc test; anti-*Crh* + mCherry^+^ vs. anti-*Crh* + mCherry^-^, *p* < 0.0001. **c.** Normalized interaction time during the repetitive social presentation test in mice expressing scrambled or anti-*Crh* shRNAs. 4 mice per group. Two-way ANOVA; trial x virus F_4,30_ = 2.00, *p* = 0.1; trial F_4,30_ = 6.07, *p* = 0.001; virus F_1,30_ = 14.62, *p* = 0.0006. **d.** Interaction time with familiar (blue) or novel (red) mouse during the recall trial of the social novelty preference test in mice expressing scrambled or anti-*Crh* shRNAs. Grey dots are different mice. Paired *t* test: *p* = 0.0095, *p* = 0.6. **e.** Discrimination indexes for social novelty preference during recall trial. Grey dots are different mice. Unpaired *t* test: *p* = 0.03. **f.** Images of mGFP expression following the injection of AAV DIO.mGFP in rdLS of *CRHR1-Cre* mice. Scale bars: 400 µm (left), 100 µm (right). **g.** C57BL/6J wild-type mice infused in rdLS with 2 µg of antalarmin dissolved in 0.6 µL of DMSO or DMSO as a control. **h.** Interaction time with familiar (blue) or novel (red) mouse during the recall trial of the social novelty preference test in mice infused with antalarmin or DMSO. Grey dots are different mice. Paired *t* tests: *p* = 0.05, *p* = 0.3. **i.** Discrimination index for social novelty preference during recall trial. Grey dots are different mice. Unpaired *t* test: *p* = 0.01. **j.** *CRH-Cre* mice injected in ILA with AAV2/2 CAG.FLEX.ArchT-TdTomato or control AAV2/2 CAG.FLEX.TdTomato. Optical ferrule implant is above rdLS. Scale bar: 500 µm. **k.** Interaction time with familiar (blue) or novel (red) mouse during the recall trial of the social novelty preference test in the same mice. Laser was on during the learning or recall trial. Grey dots are different mice. Paired *t* test: *p* = 0.045, *p* = 0.009, *p* = 0.04 and *p* = 0.8. **l.** Discrimination index for social novelty preference during recall trial of the social novelty preference test in the same mice. Grey dots are different mice. One-way ANOVA: F_3,27_ = 3.61, *p* = 0.03. Dunnett’s multiple comparisons tests: mC + light on learning vs. Arch + light on recall, *p* = 0.1; Arch + light on learning vs. Arch + light on recall, *p* = 0.03; mC + light on recall vs. Arch + light on recall, *p* = 0.04. **m.** Normalized interaction time during the repetitive social presentation test in the same mice. The laser was on during trials 1-4 of the Arch-light and mC-light groups (4 mice and 3 mice respectively). Laser was not on for the Arch-no light group (4 mice). Two-way ANOVA: trial x group F_8,40_ = 2.50, *p* = 0.03; trial F_4,40_ = 19.06, *p* < 0.0001; group F_2,40_ = 27.95, *p* < 0.0001. For the entire figure, bar graphs represent mean ± S.E.M. Grey dots are different mice.

Next, we ran *CRH-Cre* mice expressing either shRNAs for the repetitive social presentation test. Mice expressing anti-*Crh* shRNA in ILA^CRH^ cells showed very little familiarization unlike mice expressing scrambled shRNA (Fig. 3c). As a control we also ran the repetitive object presentation test. Both groups familiarized to the object to the same extent (Extended Data Fig. 8a). We also ran mice for the social novelty preference test. Mice expressing anti-*Crh* shRNA showed no preference for the novel mouse during the recall trial (Fig. 3d). Consequently, the discrimination index of this group was null (Fig. 3e). Total exploration was not different between groups (Extended Data Fig. 8b).

We then asked whether CRH release from ILA^CRH^ in rdLS cells was necessary to mediate familiarization and social novelty preference. Since CRH receptor 1 regulates social interaction and social novelty preference^31,41^, we looked whether it was expressed in the vicinity of the ILA^CRH^ cells terminals in LS. We injected *CRHR1-Cre* mice with a Cre-dependent AAV expressing GFP and confirmed expression in the dorsal part of rdLS (Fig. 3f) in the same region we found terminals of ILA^CRH^ cells (Fig. 1e). Then, we implanted mice with a cannula in rdLS and infused them with the CRHR1 antagonist antalarmin^47^ or DMSO as a control before running them for the social novelty preference test (Fig. 3g). Mice infused with DMSO exhibited normal preference for the novel mouse whereas mice infused with antalarmin showed no novelty preference (Fig. 3h-i). Total exploration time was not different between groups (Extended Data Fig. 8c). Thus, activation of the CRHR1 receptor in rdLS is necessary for social novelty preference.

Next, we used optogenetics to silence ILA^CRH^ cell terminals in rdLS. *CRH-Cre* mice were injected in ILA with Cre-dependent AAVs expressing Archaerhodopsin tagged with mCherry (Arch) or mCherry only as a control and an optical ferrule was implanted above rdLS (Fig. 3j). Light from a 568 nm laser was applied continuously during either the learning or recall trials of the social novelty preference test. When light was applied during learning both groups exhibited normal preference for the novel mouse during the recall trial. However, when light was applied during the recall trial, mice expressing Arch but not mCherry only failed to show a preference for the novel mouse (Fig. 3k-l). All groups showed the same extent of total interaction during learning and recall indicating that the manipulation did not affect sociability (Extended Data Fig. 8d-e). We also tested the mice during the repetitive social presentation test, applying the light during trials 2-4 or without light. Arch-expressing mice with light failed to familiarize to the novel mouse unlike mCherry-expressing mice with light or Arch-expressing mice with no light (Fig. 3m). Taken together these experiments suggest that CRH release from ILA^CRH^ cells in rdLS and subsequent activation of the cells expressing CRHR1 regulate the duration of social interactions with a familiar mouse.

### CRH release from ILA during familiar interaction decrease social interaction through rdLS disinhibition

What is the effect of CRHR1 activation in rdLS? We prepared acute LS slices from C57BL/6J wild-type mice and recorded spontaneous inhibitory post-synaptic currents (IPSCs) from rdLS cells, since all LS cells are GABAergic, before applying the CRHR1 agonist stressin-1 (300 nM, Fig. 4a-b)^48^. Stressin-1 application for 15 min decreased the frequency and integrated charge of spontaneous IPSCs (Fig. 4b-e). IPSC amplitude also exhibited a trend toward a decrease (Fig. 4d). This effect was not seen when rdLS neurons were recorded for 15 min without application of the agonist (Extended Data Fig. 9a). In addition, application of stressin-1 while recording from vLS neurons had no effect (Extended Data Fig. 9b), suggesting that the agonist effect was not generalized to the entire LS, consistent with the pattern of CRHR1 expression (Fig. 3f). Taken together these in vitro results suggests that CRH release disinhibits rdLS.

**Figure 4:**
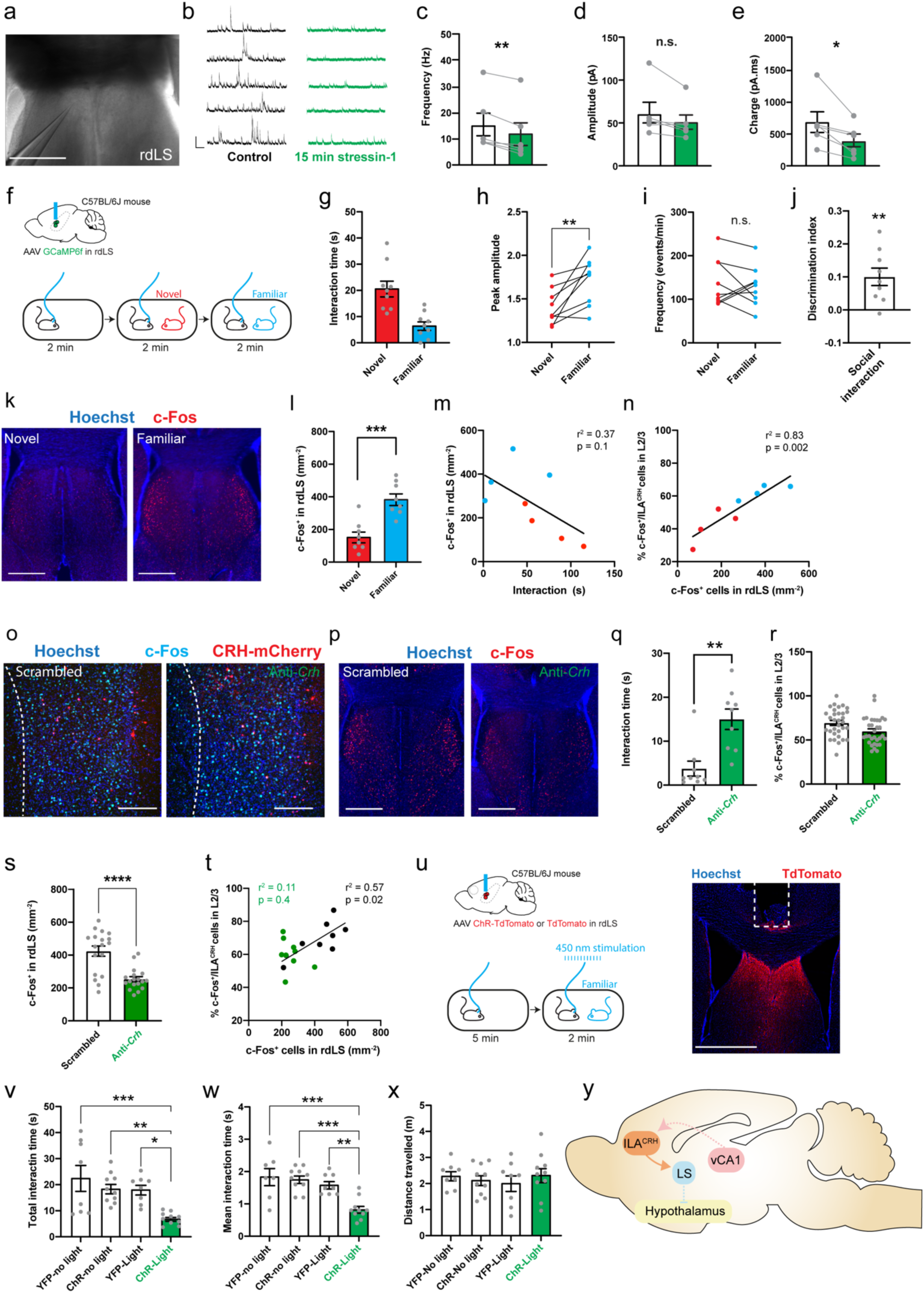
rdLS disinhibition by CRH prevents social interactions. **a.** DIC image of rdLS during patch-clamp recording. Scale bar: 500 µm. **b.** Example traces of IPSCs before or 15 min after application of 300 nM stressin-1. **c.** Frequency of IPSCs. **d.** Amplitude of IPSCs. **e.** IPSCs area under the curve. For c-e, points are obtained from individual cells recorded from separate slices in 6 mice. **f.** C57BL/6J wild-type mice injected in rdLS with AAV2/1 Syn.GCaMP6f and implanted with an optical ferrule above rdLS. Implanted mice were presented with novel and familiar mice. **g.** Interaction time during social presentation. Dots are from 9 recording sessions using 5 mice. **h.** Average peak amplitude of the z-score during presentation of a novel or familiar mouse. Paired *t* test: *p* = 0.007. **i.** Frequency of events during presentation of a novel or a familiar mouse. Paired *t* test: *p* = 0.6. **j.** Discrimination index for social familiarity preference calculated from z-scores. One-sample *t* tests compared to 0: *p* = 0.006. **k.** Immunohistochemistry images of c-Fos labelling in rdLS following social presentation with a novel or familiar mouse (same experiment than Fig. 2s). Scale bars: 500 µm. **l.** Density of rdLS cells positive for c-Fos. We made one observation on each side of a rLS section. 4 mice per group. Unpaired *t* test, *p* = 0.0003. **m.** Density of rdLS cells positive for c-Fos vs. interaction time during social interaction. Each point represents one mouse. **n.** Percentage of layer 2/3 ILA^CRH^ cells positive for c-Fos (cf. Fig. 2s) vs. density of rdLS cells positive for c-Fos following social interaction. Each point represents a mouse. **o-p.** Immunohistochemistry images of c-Fos labelling in ILA (o) and rdLS (p). *CRH-Cre* mice injected in ILA with AAV2/9 CMV-DIO-(mCherry-U6)-shRNA(anti-*Crh*) or AAV2/9 CMV-DIO-(mCherry-U6)-shRNA(scrambled) were presented with a familiar mouse for 2 min before being processed for immunohistochemistry. Scale bars: 300 µm. **q.** Duration of interaction during familiar presentation. Each point is one mouse. Unpaired *t* test, *p* = 0.001. **r.** Percentage of layer 2/3 ILA^CRH^ cells positive for c-Fos in layer 2/3 of ILA. Each point corresponds to each side of 2 sections. 9 mice per group. **s.** Density of rdLS cells positive for c-Fos. We made one observation on each side of a rLS section. 9 mice per group. Unpaired *t* test, *p* < 0.0001. **t.** Percentage of layer 2/3 ILA^CRH^ cells positive for c-Fos vs. density of rdLS cells positive for c-Fos following social interaction with a familiar mouse. Each point represents one mouse. **u.** C57BL/6J wild-type mice were injected with AA2/2 hSyn1.hChR2(H134R)-mCherry or AA2/2 hSyn1.mCherry as control and an optical fiber was implanted above the injection site. Mice were then presented to a familiar mouse for 2 min meanwhile 450 nm light was applied (20 Hz, 1 ms). Mice were also run without light as additional controls. Scale bar: 1 mm. **v.** Total interaction time with familiar mouse. Each point represents one mouse. One-way ANOVA: F_3,32_ = 7.05, *p* = 0.0009. Dunnett’s multiple comparisons tests: ChR-light vs. YFP-no light *p* = 0.0005, ChR-light vs. ChR-no light *p* = 0.006, ChR-light vs. YFP-light *p* = 0.01. **w.** Average duration of each bout of social interaction. Each point represents one mouse. One-way ANOVA: F_3,31_ = 10.62, *p* < 0.0001. Dunnett’s multiple comparisons tests: ChR-light vs. YFP-no light *p* = 0.0001, ChR-light vs. ChR-no light *p* = 0.0001, ChR-light vs. YFP-light *p* = 0.002. **x.** Total distance travelled. Each point represents one mouse. **y.** Schematic of the vCA1-ILA^CRH^-rdLS circuit. For the entire figure, bar graphs represent mean ± S.E.M.

Is rdLS disinhibited during social interactions with familiar mice and what is the effect of this disinhibition? To answer these questions, we recorded responses of rdLS neurons during interaction with a novel or familiar mouse using fiber photometry (Fig 4f). C57BL/6J wild-type mice were injected in rdLS with an AAV expressing GCaMP6f and an optical ferrule was implanted above it (Extended Data Fig. 10a-b). We presented novel and familiar mice and measured the interaction time, average peak amplitude and peak frequency (Fig. 4f-i). Presentation of a familiar mouse induced responses of larger amplitude compared to presentation of a novel mouse (Fig. 4h) despite the mice interacting less (Fig. 4g). There was no change in the frequency of events (Fig. 4i). The discrimination index based on z-score peak amplitude (see methods) showed a significant preference for the response or familiar mouse (Fig. 4j). To determine whether rdLS activity differs during novel and familiar mouse presentation, we also trained linear classifiers to discriminate between interactions with a novel or familiar mouse based on our fiberphotometry recordings. We implemented 2 classifiers using either individual recording sessions (Extended Data Fig. 10c, left) or creating a meta-session pooling all sessions (pseudo-simultaneous, Extended Data Fig. 10c, right). For each classifier, we also computed chance levels using permutation tests (grey areas). Most individual recording session yielded a decoding performance above chance with an average of 59% accuracy while the pseudo-simultaneous data yielded a decoding performance even higher (81%). This shows that rdLS can code for social familiarity. We also measure rdLS activity during the repetitive social presentation test and observed that the peak amplitude of calcium events increased from trial 1 to trial 3 when familiarization is taking place (Extended Data Fig. 10d-e). Similar to the c-Fos observation in ILA, the calcium activity anti-correlated with the amount of social interaction (Extended Data Fig. 10f).

We examined c-Fos expression in rdLS following novel or familiar encounters in the same cohort of mice than the one used to look at c-Fos expression in ILA (Fig. 2t-x). rdLS responded preferentially to familiar social presentation compared to novel social presentation (Fig. 4k-l), similar to layer 2/3 ILA^CRH^ cells response. Noteworthy, c-Fos expression was upregulated in a spatially defined band of rdLS cells bordering the lateral ventricle (Fig. 4k, right) while exposure to a novel mouse failed to activate this population (Fig. 4k, left). Taken together the fiberphotometry recordings and c-Fos labelling demonstrate that rdLS is activated preferentially during a familiar encounter compared to a novel one. As in ILA^CRH^ cells, rdLS activation correlated negatively with the amount of social interaction (Fig. 4m). We also quantified c-Fos expression in posterior dorsal LS (dLS) and posterior ventral LS (vLS) and observed no preferential response to familiar presentation compared to novel presentation nor correlation with the amount of interaction (Extended Data Fig. 11). Of note, activation of layer 2/3 ILA^CRH^ cells plotted against rdLS activation demonstrated a strong positive correlation, suggesting that one population might control the other (Fig. 4n).

Does the activation of rdLS during familiar encounters depends on CRH release from ILA^CRH^ cells? We tested this hypothesis by measuring c-Fos expression in mice where *Crh* was knocked down in ILA. *CRH-Cre* mice were injected in ILA with Cre-dependent AAVs expressing anti-*Crh* shRNA or a scrambled shRNA control, as described above. Mice were then presented with a familiar littermate and ILA and rdLS were labeled for c-Fos (Fig. 4o-p). Importantly, anti-*Crh* expressing mice interacted more with a familiar mouse than control mice, suggesting that *Crh* reduces social interaction with a familiar mouse (Fig. 4q). Loss of CRH did not alter c-Fos expression in layer 2/3 ILA^CRH^ cells (Fig. 4r) while it strongly decreased c-Fos expression in rdLS (Fig. 4s). While control mice exhibited the normal correlation between c-Fos levels in layer 2/3 ILA^CRH^ and rdLS, test mice depleted of *Crh* in ILA did not (Fig. 4t). A similar approach was used previously to disrupt memory retrieval^49^. Taken together these experiments demonstrate that CRH release from ILA during familiar encounter disinhibits a specific population of rdLS cells bordering the ventricles.

Our findings suggest that rdLS disinhibition suppresses social interactions. To explore this further, we examined the effects of optogenetic activation of rdLS neurons. We injected an AAV expressing Channelrhodopsin (ChR) tagged with mCherry or mCherry only in rdLS of C57BL/6J wild-type mice and implanted an optical ferrule above it (Fig. 4u). We then presented a familiar mouse for 2 min while stimulating ChR using a 450 nm laser (1 ms stimulation at 20 Hz) and measured the interaction time as well as the mean duration of each interaction bout. As an additional control, we measured behavior in ChR-expressing mice without laser stimulation. Activation of rdLS decreased the amount of social interaction with familiar mice (Fig. 4v) due to shorter interactions each time the mice met (Fig. 4w). Similar results were observed during novel mouse encounters (Extended Data Fig. 12). These optogenetic experiments demonstrate that rdLS is able to decrease social interactions. Altogether our study shows that ILA^CRH^ cells are activated during familiar mouse encounters, leading to release of CRH in rdLS causing its disinhibition (Fig. 4x). Disinhibition of rdLS in turn prevents social interaction which explain the decreased social interaction observed during familiarization. Inhibition of social interaction with familiar mice promotes social novelty preference.

### Emergence of the ILA^CRH^ cells to rdLS circuit is responsible for the shift in social preference observed in young mice

In contrast to adults, young rodents prefer to interact with their familiar siblings compared to novel pups^3,4,50^. We tested when the shift in social preference occurs in. mice by giving young C57BL/6J wild-type mice the choice to interact with their familiar siblings or with unfamiliar non-siblings every day from P7 to P21. Between P7 and P15, mice preferred to interact with their siblings (Fig. 5a) and their discrimination index was strongly skewed toward familiarity (Fig. 5b). Preference then gradually shifted toward novel mice (Fig. 5a) and the index leaned toward social novelty after P16 (Fig. 5b).

**Figure 5:**
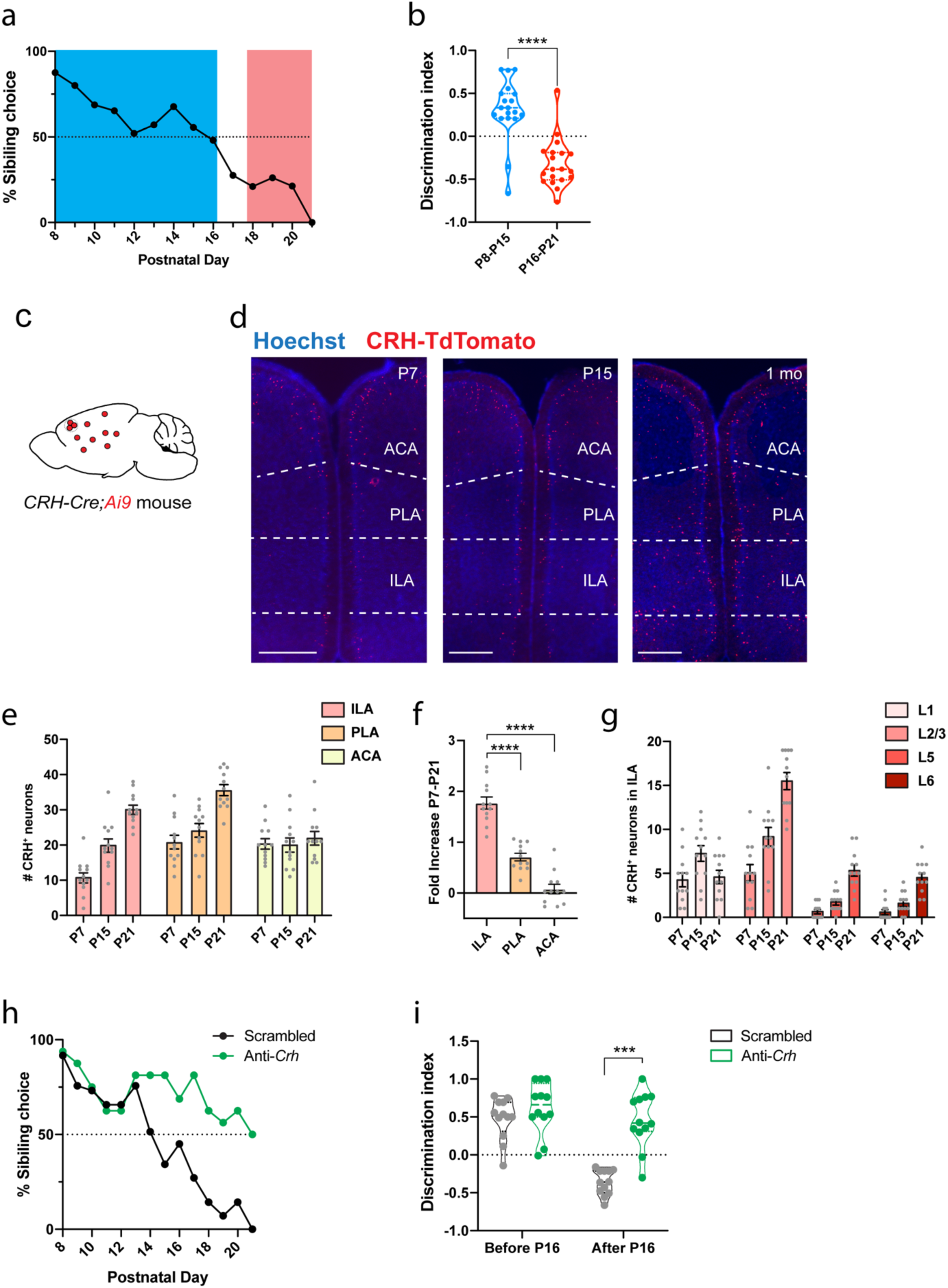
IL^CRH^ to LS circuit emerges during development to control kinship preference. **a.** Percentage of familiar choice during development, 19 mice. **b.** Discrimination index for familiar kin before and after postnatal day 16. Each point represents a mouse, 19 mice. Unpaired *t* test, *p* < 0.001. **c.** *CRH-Cre;Ai9* mice. **d.** mPFC images of *CRH-Cre;Ai9* mice at P7, P15 or P21. Scale bars: 500 µm. **e.** Number of CRH^+^ cells in ILA, PLA and ACA during development. Each point represents one observation made on each side of 2 section, 3 mice per group. **f.** Fold-increase of CHR^+^ cells between P7 and P21. P21 values compared to the average P7 value. One-way ANOVA: F_2,33_ = 75.68, *p* < 0.0001. Dunnett’s multiple comparisons tests: ILA vs. PLA, *p* < 0.0001 and ILA vs. ACA, *p* < 0.0001. **g.** Number of CRH^+^ cells per ILA layers during development. Each point represents one observation made on each side of 2 section, 3 mice per group. **h.** Percentage of familiar choice during development in *CRH-Cre* mice injected in ILA with AAV2/9 CMV-DIO-(mCherry-U6)-shRNA(anti-*Crh*) to downregulate *Crh* or control AAV2/9 CMV-DIO-(mCherry-U6)-shRNA(scrambled). 12 pups per group. Chi-square test: *p* < 0.0001. **i.** Discrimination index for familiar kin before and after postnatal day 16. Each point represents a mouse, 12 pups per group. Unpaired *t* tests: *p* = 0.3 and *p* < 0.0001. For the entire figure, bar graphs represent mean ± S.E.M.

Both the PFC and LS have been shown to control preference for a novel conspecific compared to a familiar one and the LS is also involved in the preference young rats display for their own (familiar) kin compared to non-kin^3^. We then asked when the ILA cells begin to express CRH and whether the emergence of the ILA to rdLS circuit contributes to the shift in social preference. We counted CRH-TdTomato^+^ cells at P7, P15 and P21 in *CRH-Cre;Ai9* mice (Fig. 5c-d) and observed a strong increase in the density of ILA^CRH^ cells from P7 to P21 (Fig. 5e). Interestingly, the increase was the strongest in ILA compared to other prefrontal regions (Fig. 5f). Within ILA, the increase of CRH^+^ cells proceeded from an increase in CRH^+^ cells located in layer 2/3 (Fig. 5g), which is the layer containing ILA^CRH^ cells projecting to rdLS (Fig. 1). Closer inspection of CRH^+^ cells in PLA and ACA revealed less or no increase in layer 2/3 (Extended Data Fig. 13). Overall, these experiments demonstrate that the ILA^CRH^ to rdLS circuit is strengthened during the shift from social familiarity to social novelty preference.

We next probed whether the emergence of the circuit causes the shift in preference. P5 pups were injected with AAVs expressing anti-*Crh* or scrambled shRNAs. We reasoned the AAVs would be taken up in many ILA neurons so that, as soon as CRH expression begins, so would the Cre recombinase expression and therefore shRNA expression under the control of a fast-expressing U6 promoter. 2 days after the injection, we began testing the injected pups for social preference and observed a preference shift at P14 in the control group similar to our previous experiment on wild-type mice (Fig. 5a). The test group lacking *Crh* however continued to exhibit familiar preference until P20 (Fig. 5h). Consistently, discrimination indexes for the control group inverted before and after P16 showing the shift in social preference while the ones for the *Crh*-depleted group remained oriented toward familiar choice (Fig. 5i). Overall, these experiments demonstrate that increased CRH expression in ILA is responsible for the shift in social preference displayed by young mice.

## Discussion

We show that ILA^CRH^ cells respond to social interaction with familiar over novel mice and release CRH into rdLS in order to suppress social interactions with familiar mice through LS disinhibition. During familiarization, increasingly responsive ILA^CRH^ cells control the decrease in interaction as a novel mouse becomes familiar. When given the choice between a familiar and a novel mouse, this circuit suppress interaction with the familiar which promote SNP.

We asked before whether ILA^CRH^ cells control social memory or rather downstream processes, including SNP that utilize social memory cues? Silencing ILA^CRH^ cell terminals to rdLS during the recall trial but not during the learning trial disrupts SNP (Fig. 3k-l), showing that ILA^CRH^ cells are not contributing to social memory formation. This is similar to previous work showing CRH-binding protein being important for the recall but not learning phase of social recognition^31^. ILA^CRH^ cells could however still be involved in the social memory recall. Knocking down *Crh* in ILA increases social interactions with familiar mice (Fig. 4r) while keeping the c-Fos expression in ILA unchanged. This demonstrate that *Crh* is not needed for ILA^CRH^ activation by familiarity cues while it is sufficient to regulate social interactions with a familiar animal.

Previous hypotheses about the mechanisms underlying SNP supposed the existence of a circuit promoting interaction with novel mice, perhaps under control of the rewarding properties of social novelty. In addition, the kin (mother or siblings) preference^18,50^ displayed by young mice supposes the existence of other circuits controlling social preference. Very little is known however about the mechanisms supporting the rewarding properties of social cues. The lateral habenula, nucleus accumbens, dorsal raphe nucleus and ventral tegmental area modulate social reward^51–55^, some of them under control of oxytocin^51–53^. Subsequent studies should aim to characterize how social novelty reward facilitate interaction with novel mice to regulate social preference. The lateral septum, which is heavily modulated by dopamine, vasopressin and oxytocin^19^, may act as a hub and also integrate inputs promoting interaction with novel mice in order to regulate social preference.

How specific is the regulation of social preference by ILA^CRH^ cells? We demonstrate that ILA^CRH^ cells control memory-based social preference but not object preference. Whether ILA^CRH^ cells control other social preferences such as preferences based on sex, strain, kinship or anxiety (mice prefer to interact with non-stressed mice)^13^ remain to be determined. Overall, ILA^CRH^ cells integrate information from many brain regions which are likely to provide various social cues about nature and identity of the stimulus animals and our study supposes that social cues of negative valence activate excitatory neurons projecting on ILA^CRH^ cells. Social memories of previous encounters are known to be stored in the pyramidal neurons of the ventral CA1 region of the hippocampus^56^ and vCA1 projection to the mPFC is necessary for behaviors relying on social memory cues^57,58^. We demonstrate that ILA^CRH^ cells receive inputs from vCA1. However, whether vCA1 neurons projecting to ILA^CRH^ cells carry social familiarly information remains to be investigated. Furthermore, olfactory cues are known to guide sex- and hierarchy-based preference^9,59^ and ILA^CRH^ cells receives inputs from the orbitofrontal cortex which is known to integrate olfactory cues^60,61^. Interestingly, ILA^CRH^ cells also receive direct feedback from dLS which is striking since most LS projections descends to subcortical areas. This is similar to previous findings reporting LS^SST^ cells project to the ILA and to a lesser extent to the PLA^62^. Finally, ILA^CRH^ cells also receive inputs from many local PFC neurons which are likely to regulate ILA^CRH^ cells and therefore participate in the decision to interact.

Young mice display kin preference (for mother and siblings) during the first weeks of life^3,18^. Here, we show how young mice reliably display social preference toward their siblings versus age-matched pups until CRH increase triggers a shift in preference toward the normotopic adult behavior. While defenseless pups^63^ need to rely on the safety of their nest and company of their siblings, older and more able young mice^63^ must leave their kin and venture out of the nest in order to sample resources (feeding behavior) and interact with novel conspecifics (reproductive behavior). Interestingly, humans can suffer from social separation anxiety disorder, which manifests itself as an “unusually strong fear or anxiety to separating from people they feel a strong attachment to”^64^. Patients present unusual distress at the discussion or experience of being parted from their attachment figure and a refusal to leave the attachment figure. In addition, CRH has been involved in various anxiety disorders, including social phobia^34,65^. Similar to our findings, familiarity cues activate the human PFC^66,67^ and septal^68^ regions, supporting the idea that the circuit we described in the mouse is conserved in humans. One of the causes for social separation anxiety disorder may be that patients exhibit low CRH level in the PFC, preventing them from seeking social novelty.

## Author Contributions

Conceptualization: N.S.L.R., J.S., A.A and F.L.; Investigation, F.L.; In vitro intra-cellular recordings: N.S.L.R.; Behavioral assays and viral injections: N.S.L.R., P.S.D. and F.L. and O.M.L.; Immunohistochemistry and in situ hybridization: N.S.L.R., A.R, Y.N. and F.L.; shRNA design C.S., Fiberphotometry: N.S.L.R., Y.N.; Classifier analysis: R.N.; Writing – Original draft: F.L.; Writing – Review and editing: N.S.L.R. and F.L.; Visualization: N.S.L.R. and F.L.; Supervision: F.L.; Funding acquisition: F.L.

## Acknowledgments

We thank Antoine Besnard, Claudio Elgueta, Azahara Oliva, Isabel Otaño-Perez and Steve Siegelbaum for critical reading of the manuscript and insightful comments. We also thank Christina Fregola and Georg Keller for providing viral vectors. Finally, we thank Jan Deussing for sharing the *CRHR1-Cre* mouse line. F.L. acknowledges support from an ERC starting investigator grant (H202O-ERC-STG/0784-Ref.949652), a CIDEGENT grant from the Valencian Community and a NARSAD young investigator grant from the Brain and Behavior research foundation, funded by the Osterhaus Family. R.N. is supported by NSF Neuronex (1707398) and the Gatsby Charitable Foundation (GAT3708).

---

**Extended Data Figure 1:**
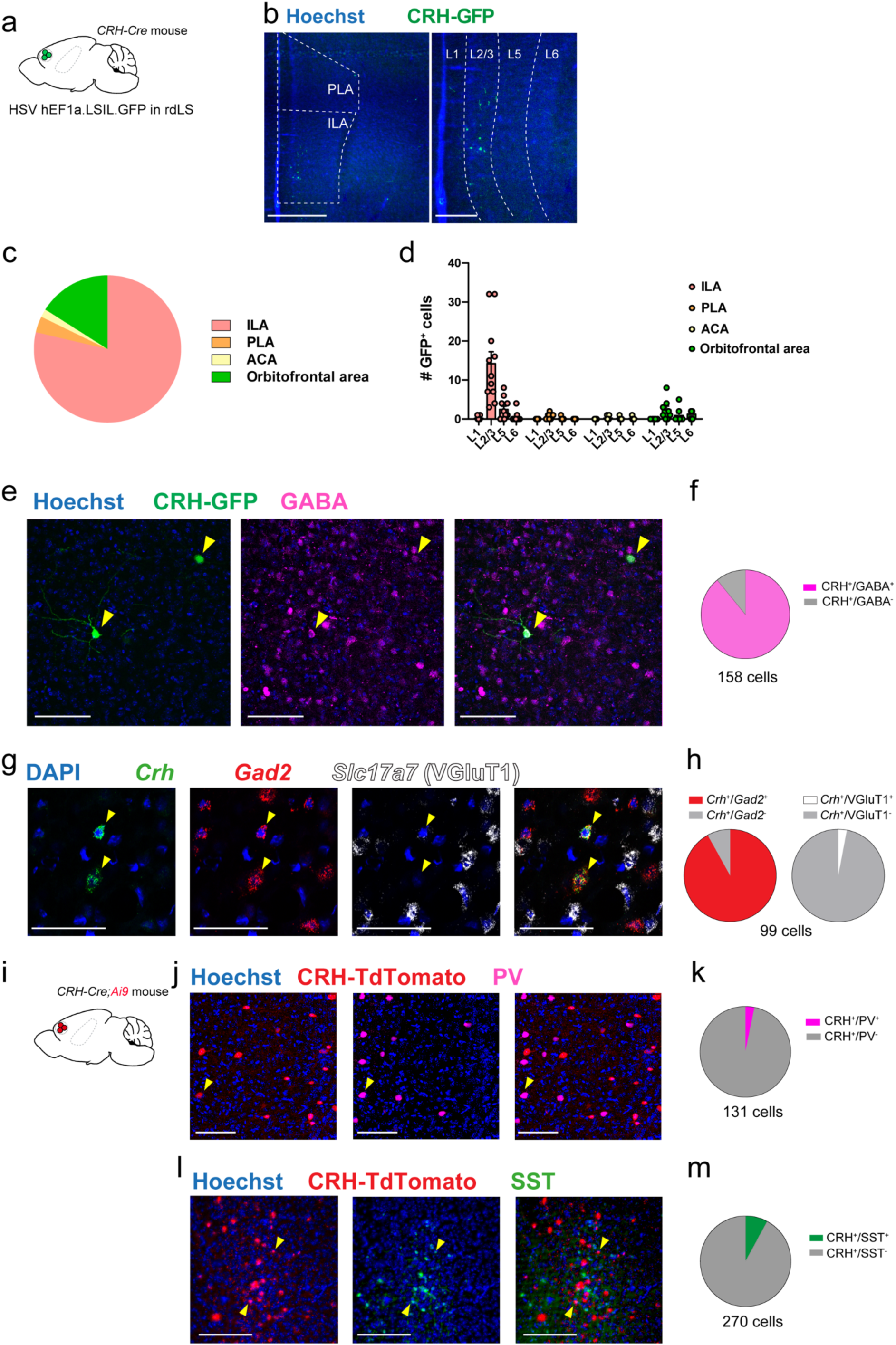
Molecular identity of ILA^CRH^ cells. **a.** *CRH-Cre* mice injected with HSV hEF1a.LSIL.GFP in rdLS. N = 3 mice. **b.** Images of the mPFC and ILA showing GFP^+^ cells retrogradely labelled. Scale bars: 500 µm (left) and 200 µm (right). **c.** Distribution of GFP^+^ cells per regions. **d.** Number of GFP^+^ cells per region and layers. **e.** Immunohistochemistry images of the ILA labelled for GABA. Scale bars: 100 µm. **f.** Associated distribution of GFP^+^ cells. **g.** In situ hybridization images of the ILA labeled for *Crh*, *Gad2* and *Slc17a7* (VGluT1). N = 3 mice. Scale bars: 50 µm. **h.** Associated distribution of *Crh*^+^ cells. **i.** *CRH-Cre;Ai9* mice. **j.** Immunohistochemistry images of the ILA labelled for PV. N = 3 mice. Scale bars: 100 µm. **k.** Associated distribution of TdTomato^+^ cells. **l.** Immunohistochemistry images of the ILA labelled for SST. N = 3 mice. Scale bars: 200 µm. **m.** Associated distribution of TdTomato^+^ cells.

**Extended Data Figure 2:**
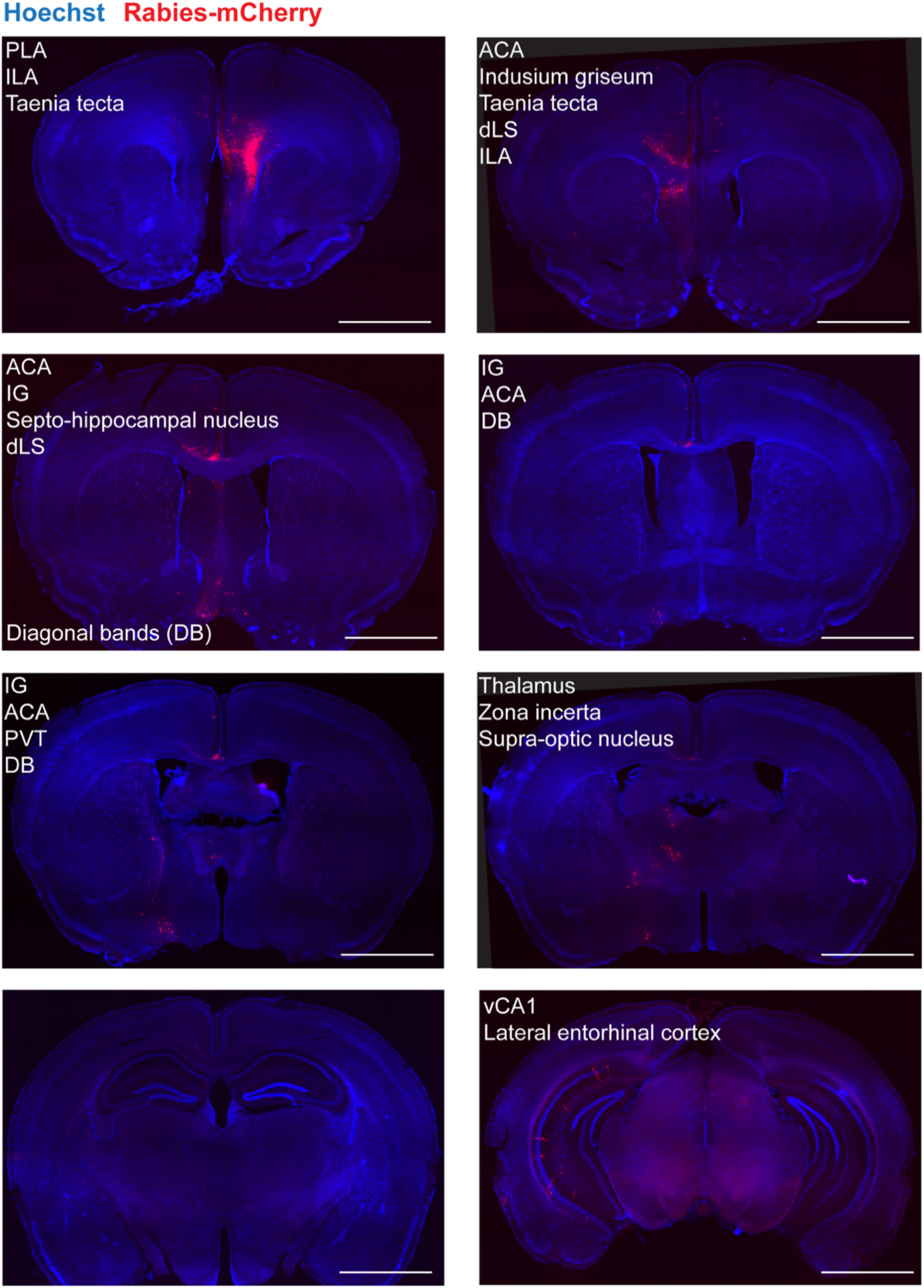
Immunohistochemistry images following rabies-mCherry expression in ILA^CRH^ cells. PLA: peri-limbic area; ACA: anterior cingulate area; IG: Indusium griseum; SHi: septo-hippocampal nucleus; rdLS: rostro-dorsal lateral septum; DB: diagonal bands; PVT: paraventricular nucleus of the thalamus. Scale bars: 2 mm.

**Extended Data Figure 3:**
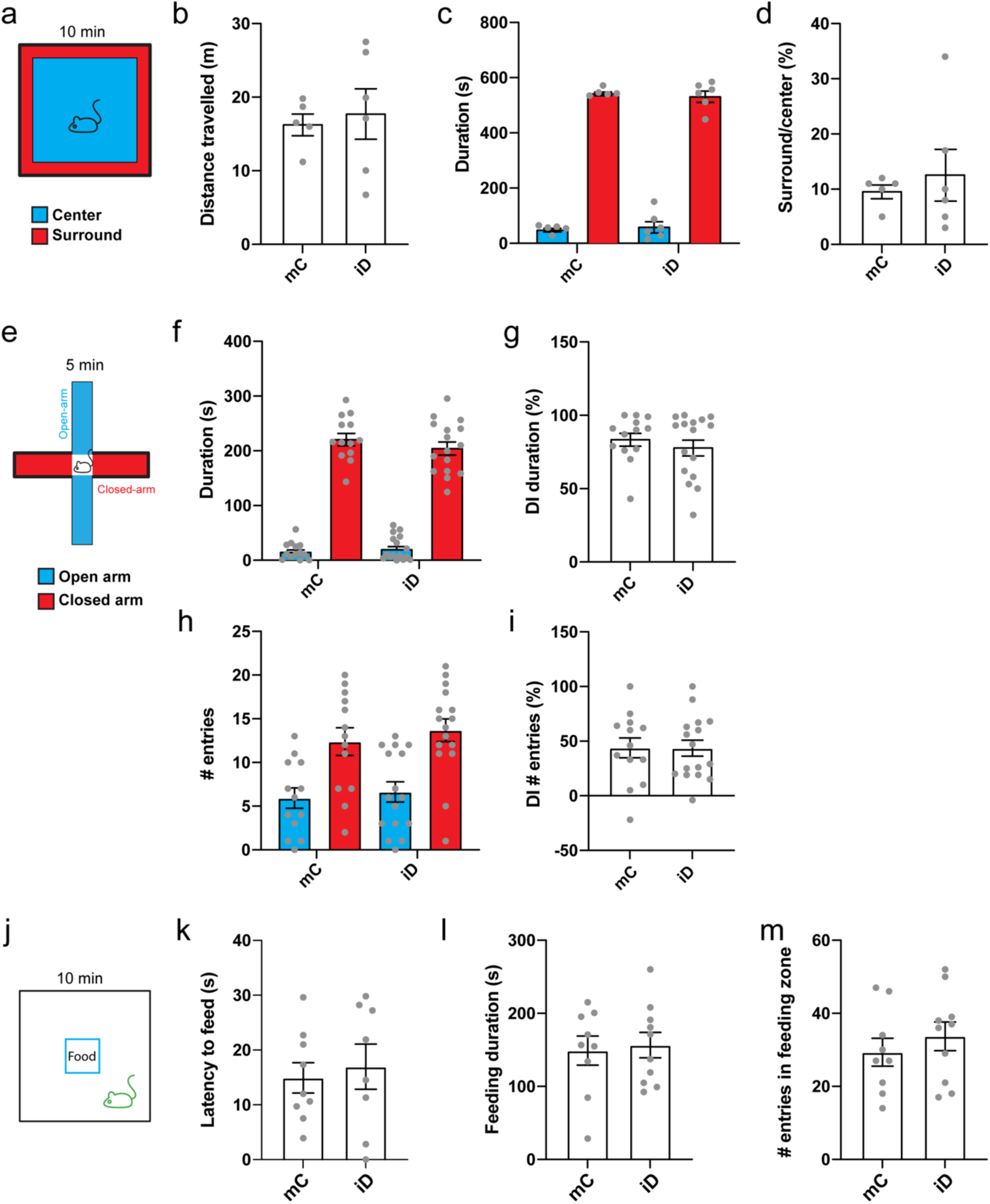
Behavioral controls for chemogenetic silencing of ILA^CRH^ cells. *CRH-Cre* mice injected in ILA with AAV2/8 hSyn.DIO.hM4D(Gi)-mCherry (iD) or AAV2/8 hSyn.DIO.mCherry (mC). **a.** Schematic of the open-arena test. **b.** Total distance travelled during open-field test. **c.** Time spent in the center or surround of the open-field. **d.** Ratio of the time spent in the center/surround. **e.** Schematic of the elevated-plus maze test. **f.** Time spent in the open or closed arms. **g.** Discrimination indexes for closed arm preference using the time spent in the arms. **h.** Number of entries in the open or closed arms. **i.** Discrimination indexes for closed arm preference using the number of arm entries. **j.** Schematic of the novelty suppressed feeding test. **k.** Latency to feed. **l.** Feeding duration. **m.** Number of entries in the feeding zone. For the entire figure, bar graphs represent mean ± S.E.M. Grey dots are different mice.

**Extended Data Figure 4:**
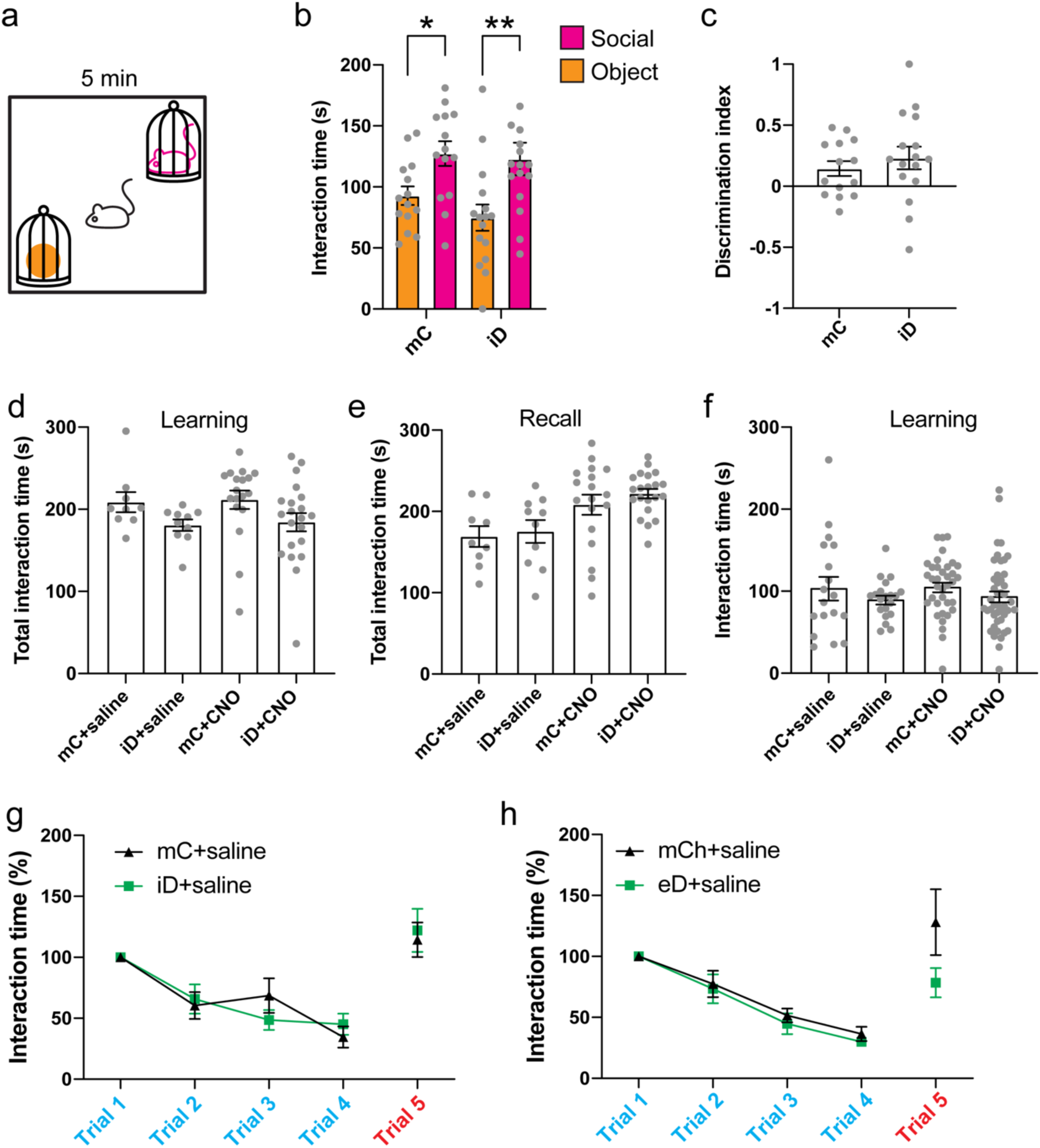
Social behavior controls for chemogenetic silencing of ILA^CRH^ cells. *CRH-Cre* mice injected in ILA with AAV2/8 hSyn.DIO.hM4D(Gi)-mCherry (iD) or AAV2/8 hSyn.DIO.mCherry (mC). **a.** Schematic of the sociability test. **b.** Interaction times with mouse (purple) or object (orange). Paired *t* tests: *p* = 0.01, *p* = 0.009. **c.** Discrimination indexes for social preference. One-sample *t* tests compared to 0: *p* = 0.03 and *p* = 0.02. **d-e.** Total interaction times during learning (d) or recall trial of the social novelty preference test. **f.** Interaction time with each novel mouse during the learning trial of the social novelty preference test. One-way ANOVA, F_3,114_ = 0.8694, *p* = 0.5. **g.** Normalized interaction times during social presentations (inhibitory DREADD-expressing mice and controls injected with saline). 8 mice per group. **h.** Normalized interaction times during social presentations (excitatory DREADD-expressing mice and controls injected with saline). 8 mice per group. For the entire figure, bar graphs represent mean ± S.E.M. Grey dots are different mice expect for f. where two observation per mice could be made since the mice interacted with two novel mice during the learning trial.

**Extended Data Figure 5:**
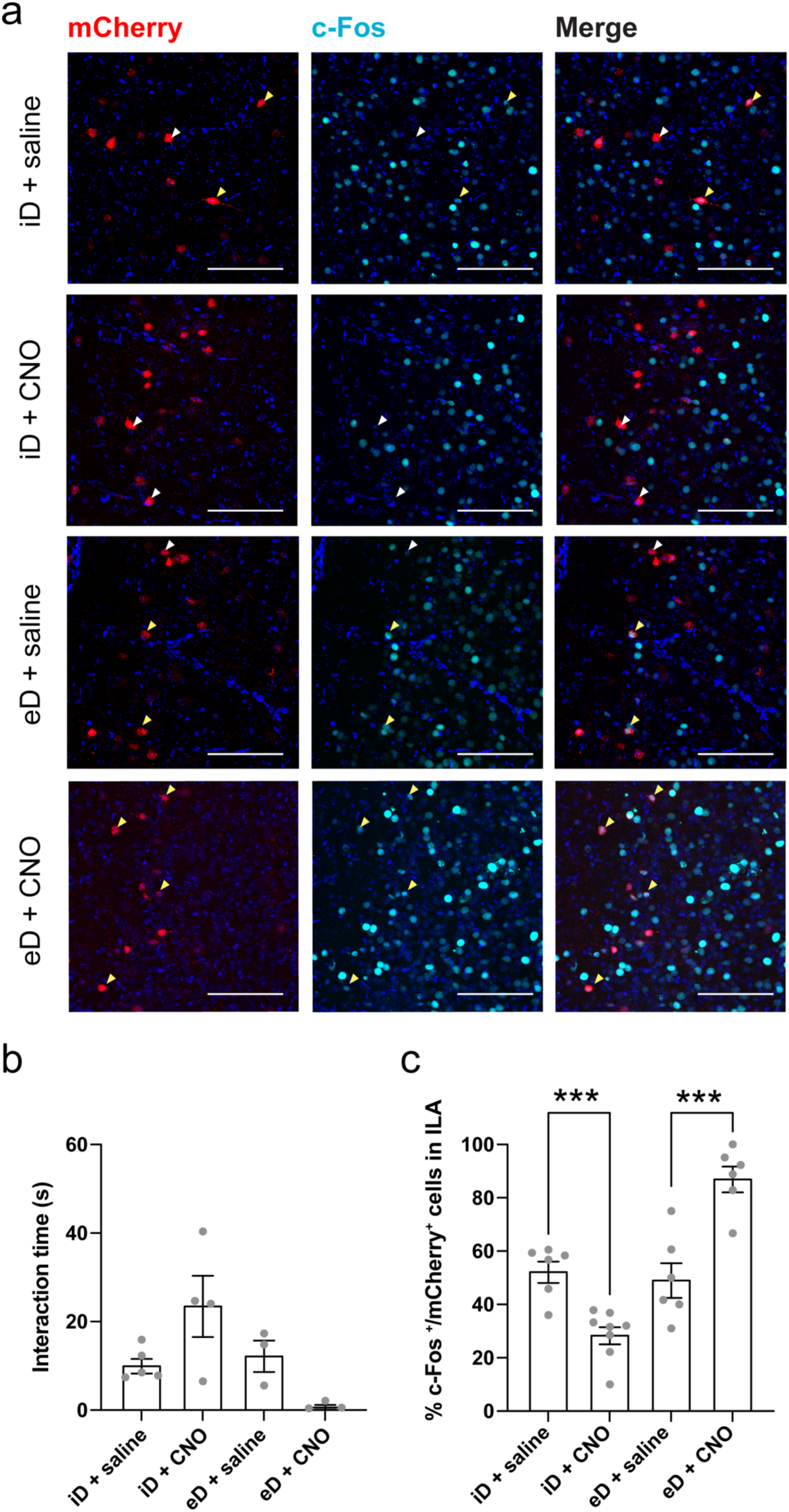
DREADD modulate ILA^CRH^ cells activity. *CRH-Cre* mice injected in ILA with AAV2/8 hSyn.DIO.hM4D(Gi)-mCherry (iD) or AAV5 hSyn.DIO.hM3D(Gq)-mCherry (excitatory DREADD). Mice received CNO or saline i.p. injection 30 min before being presented to a familiar animal for 2 min. Mice were thereafter perfused and processed for c-Fos labelling. Scale bars: 100 µm. **a.** Immunohistochemistry images of the ILA labelled for c-Fos. **b.** Interaction times. Each point is one mouse. **c.** Percentage of mCherry^+^ cells in ILA expressing c-Fos. Unpaired *t* tests: *p* = 0.0005 and *p* = 0.0009.

**Extended Data Figure 6:**
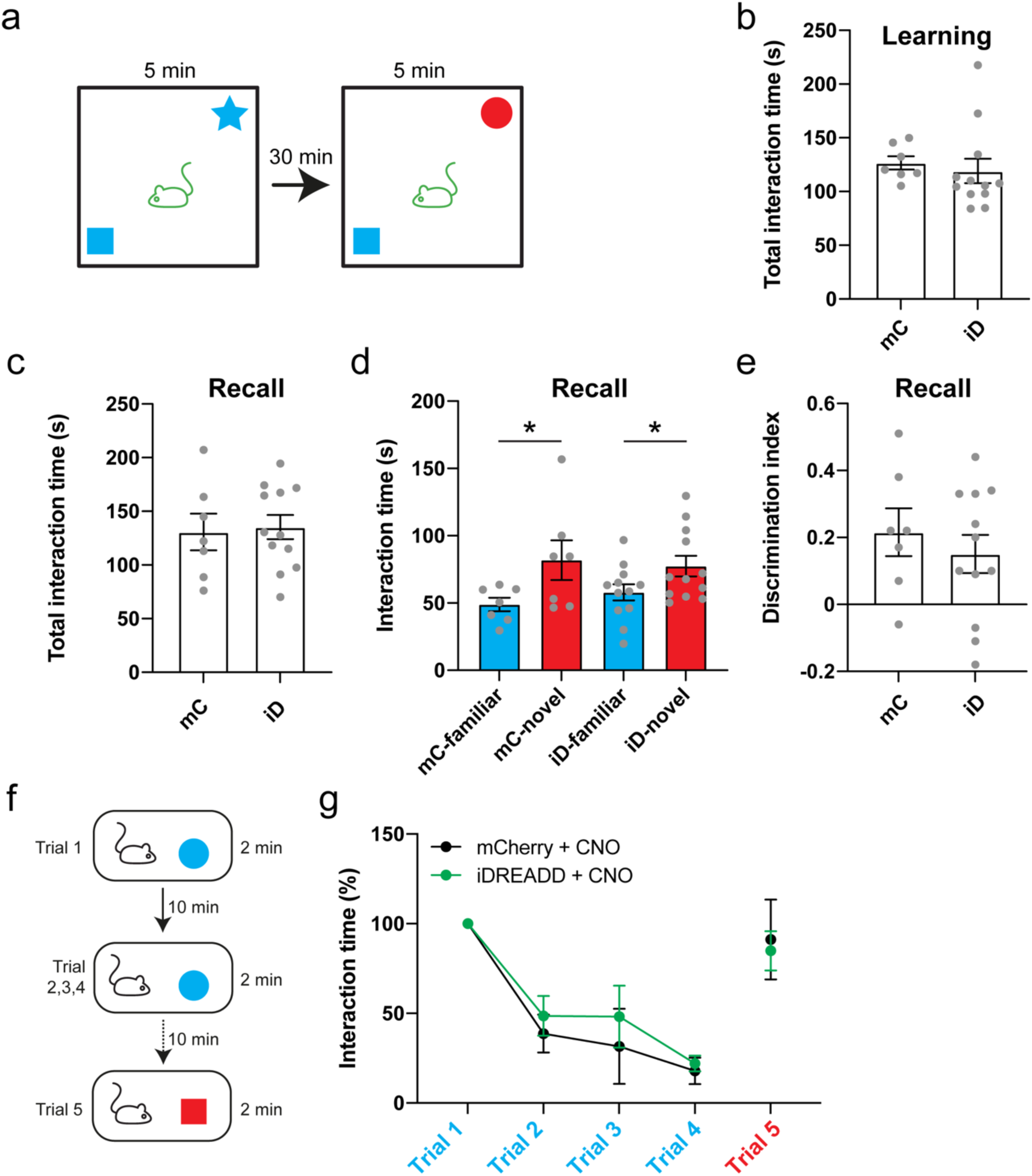
Object memory controls for chemogenetic silencing of ILA^CRH^ cells. *CRH-Cre* mice injected in ILA with AAV2/8 hSyn.DIO.hM4D(Gi)-mCherry (iD) or AAV2/8 hSyn.DIO.mCherry (mC). **a.** Schematic of the object recognition memory test. **b-c.** Total interaction times during learning (d) or recall trial of the object memory test. Grey dots are different mice. **d.** Interaction times with familiar (blue) or novel object (red). Grey dots are different mice. Paired *t* tests: *p* = 0.05 and *p* = 0.03. **e.** Discrimination indexes for novel object preference. Grey dots are different mice. One-sample *t* tests compared to 0: *p* = 0.02 and *p* = 0.02. Unpaired *t* test between groups: *p* = 0.5. **f.** Schematic of the repetitive object presentation test. **g.** Normalized interaction times during the repetitive object presentation test (inhibitory DREADD-expressing mice and control mice injected with 0.5 mg/kg CNO). 4 mice per group, 8 mice total. Two-way ANOVA: trial x virus F_4,30_ = 0.233, *p* = 0.91; trial F_4,30_ = 14.02, *p* < 0.0001; virus F_3,40_= 0.36, *p* = 0.56. For the entire figure, bar graphs represent mean ± S.E.M.

**Extended Data Figure 7:**
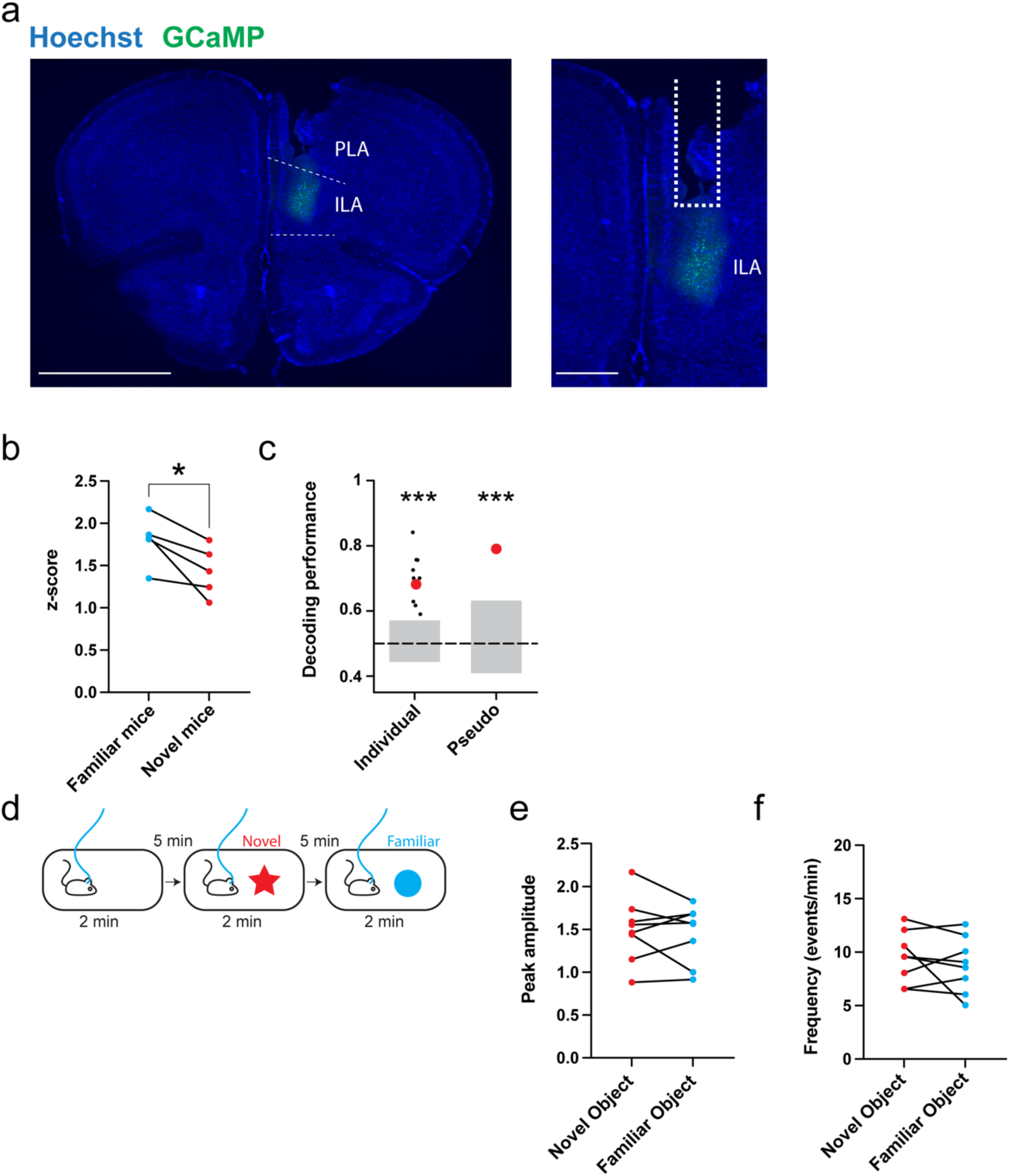
Fiber-photometry controls. **a.** Immunohistochemistry images of GCaMP expression in the ILA *of CRH-Cre* mice. Scale bars: 2 mm (left) and 200 µm (right). **b.** Average peak amplitude of the z-score during presentation of familiar then novel mouse. Paired *t* test, *p* = 0.03. **c.** Decoding performance for familiarity versus novelty from individual recordings or pseudo-simultaneous data. Small black dots on the left are the results from individual recording sessions using 20 cross-validation iterations, large red dot is the average. Red dot on the right is the result of pseudo-population analysis from 100 cross-validation iterations. Grey areas denote chance level computed using permutation tests (2.5 – 97.5 percentiles in distribution of shuffled decoding performances). In both cases, statistical significance is determined by the probability of drawing the observed decoding performance from the distribution of shuffled decoding performances (null-hypothesis). *p* < 0.001 (two-tailed permutation test, see Methods). **d.** Schematic of the object presentations. **e.** Average peak amplitude of the z-score during presentation of a novel or familiar object. Dots are different sessions using 3 mice. **f.** Frequency of calcium events during presentation of a novel or familiar object. Dots are different sessions using 3 mice.

**Extended Data Figure 8:**
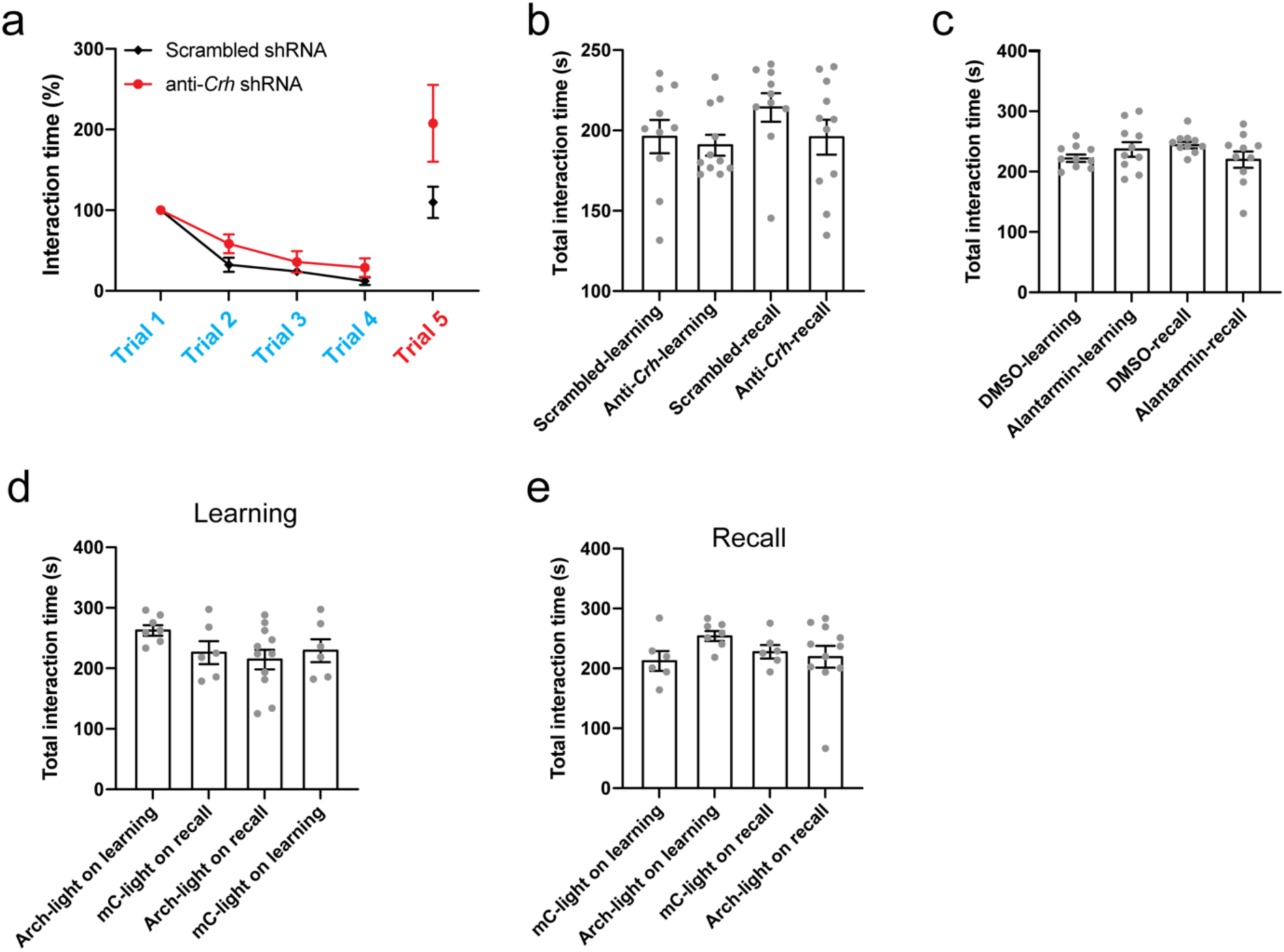
Social behavior controls for CRH release from IL to PFC. **a.** Normalized interaction times during repetitive object presentation following expression of a shRNA against *Crh* or a scrambled shRNA in ILA^CRH^ cells. **b.** Total interaction time during learning and recall of the social novelty preference test in the same mice. **c.** Total interaction time during learning and recall of the social novelty preference test following antalarmin infusion. **d-e.** Total interaction time during learning and recall of the social novelty preference test following Arch or mCherry expression in ILA^CRH^ and silencing of their terminals in LS. For the entire figure, bar graphs represent mean ± S.E.M. Grey dots are different mice.

**Extended Data Figure 9:**
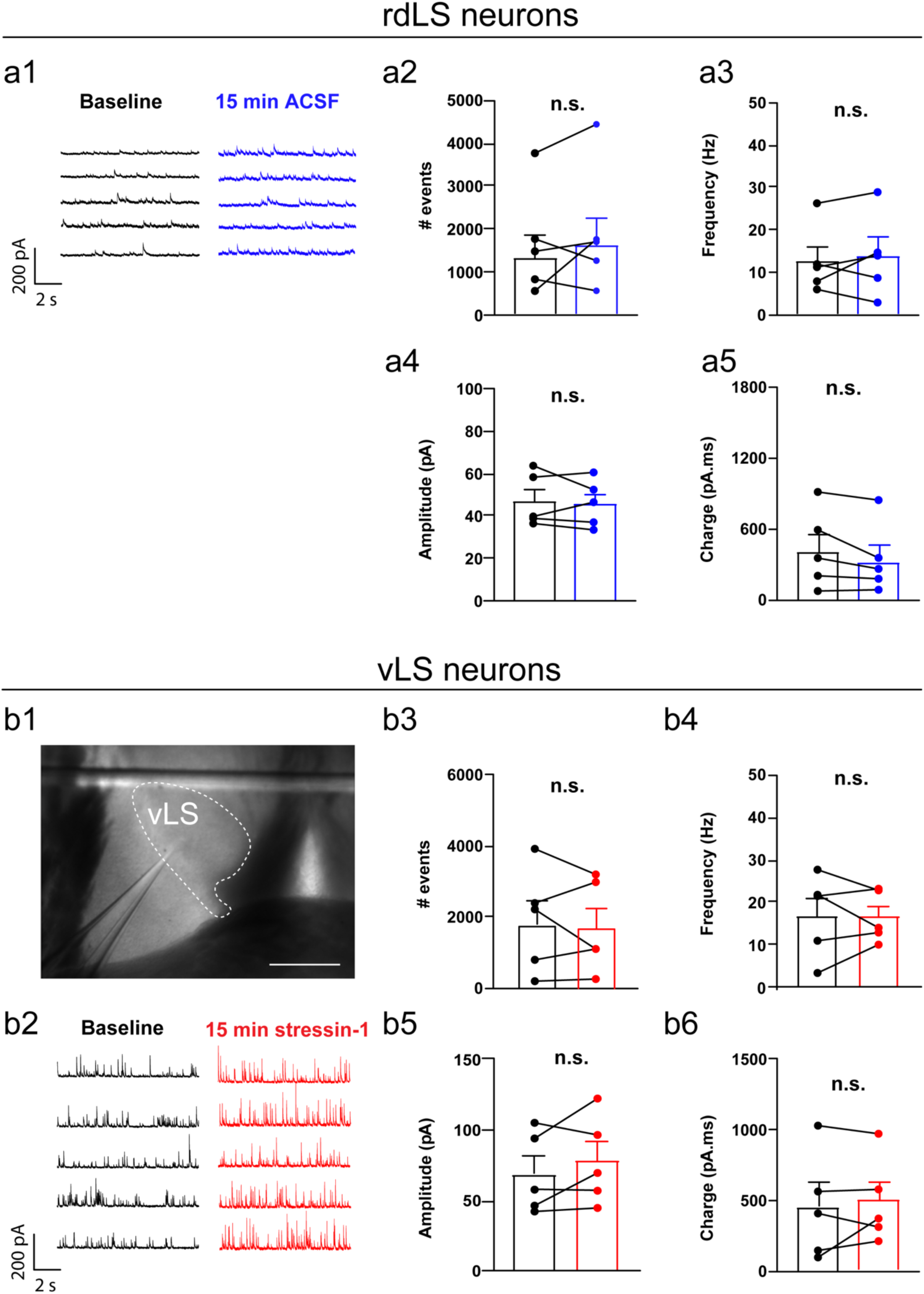
controls for in vitro electrophysiology recordings in LS. **a.** In vitro whole-cell patch-clamp of rdLS neurons. **a1.** Example trace of IPSCs before or 15 min after application of ACSF. **a2.** Number of IPSCs. Points are individual cells recorded in 5 mice. **a3.** Frequency of IPSCs. **a4.** Amplitude of IPSCs. **a5.** IPSCs area under the curve. **b.** Neurons recorded in vLS before and after application of 300 nM stressin-1. **b1.** DIC image of the LS region where cells were recorded. Scale bar: 200 µm. **b2.** Example trace of IPSCs before or after 15 min 300 nM stressin-1. **b3.** Number of IPSCs. Points are individual cells recorded in 5 mice. **b4.** Frequency of IPSCs. **b5.** Amplitude of IPSCs. **b6.** IPSCs area under the curve. For the entire figure, bar graphs represent mean ± S.E.M.

**Extended Data Figure 10:**
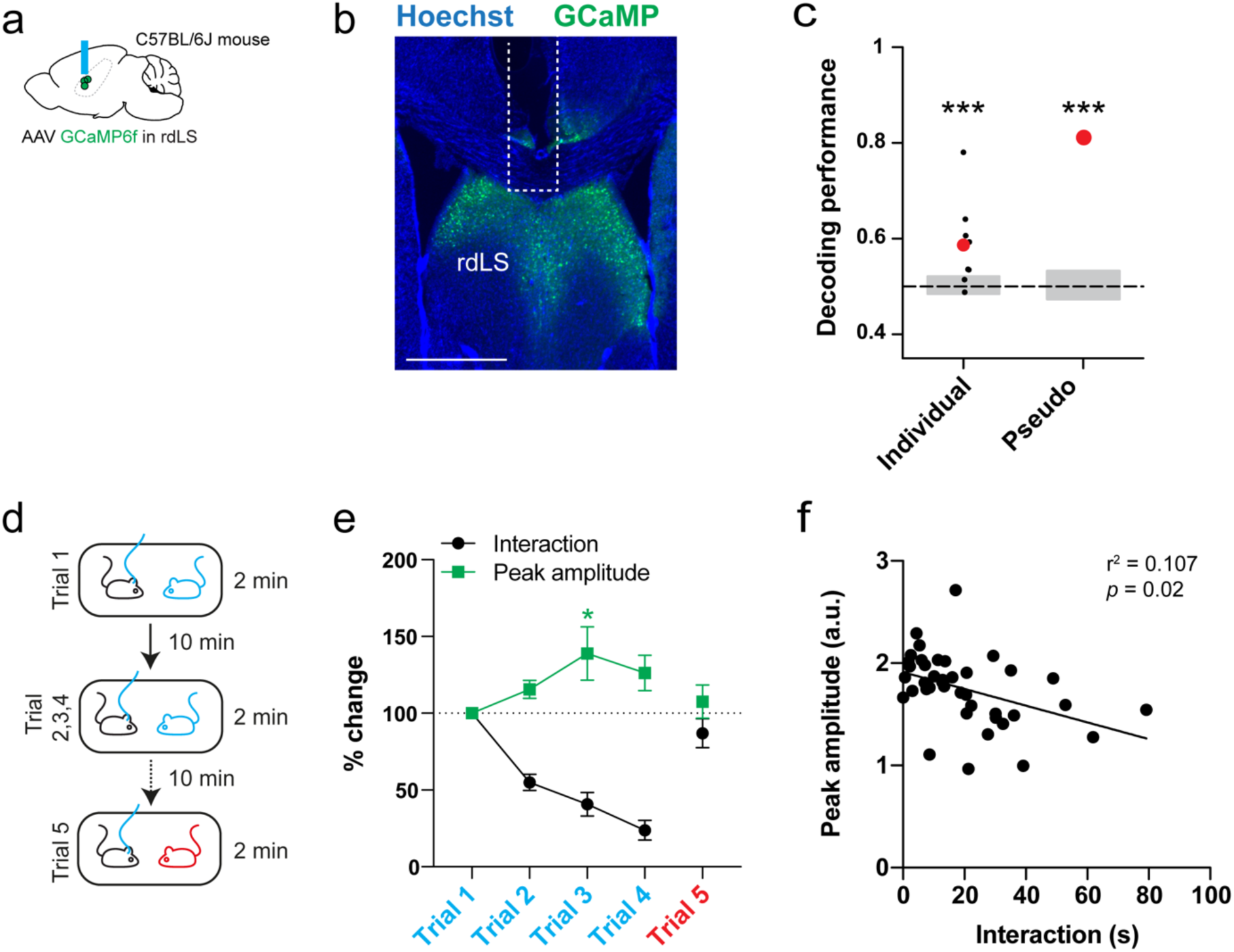
Fiber-photometry in rdLS during the repetitive social presentation test. **a.** C57BL/6J wild-type mice injected in rdLS with AAV2/1 Syn.GCaMP6f and implanted with an optical ferrule above rdLS. **b.** Immunohistochemistry image showing GCaMP expression in rdLS. Scale bar: 500 µm. **c.** Decoding performance for familiarity versus novelty from individual recordings or pseudo-simultaneous data. Small black dots on the left are the results from each individual recording sessions using 20 cross-validation iterations, large red dot is the average. Red dot on the right is the result of pseudo-population analysis from 100 cross-validation iterations. Grey areas denote chance level computed using permutation tests (2.5 – 97.5 percentiles in distribution of shuffled decoding performances). In both cases, statistical significance is determined by the probability of drawing the observed decoding performance from the distribution of shuffled decoding performances (null-hypothesis). *p* < 0.001 (two-tailed permutation test, see Methods) **d.** Schematic of the repetitive social presentation test. **e.** Fiber-photometry recording during repetitive social presentation test (10 recording sessions in 5 mice). One-way ANOVA (trial 1-4) F_3,27_ = 3.389, *p* = 0.03 followed by Dunnett’s multiple comparison tests compared to trial 1: trial 2 *p* = 0.5, trial 3 *p* = 0.01 and trial 4 *p* = 0.1. **f.** Interaction times (trial 1 to trial 4) vs. z-scores during the same experiment.

**Extended Data Figure 11:**
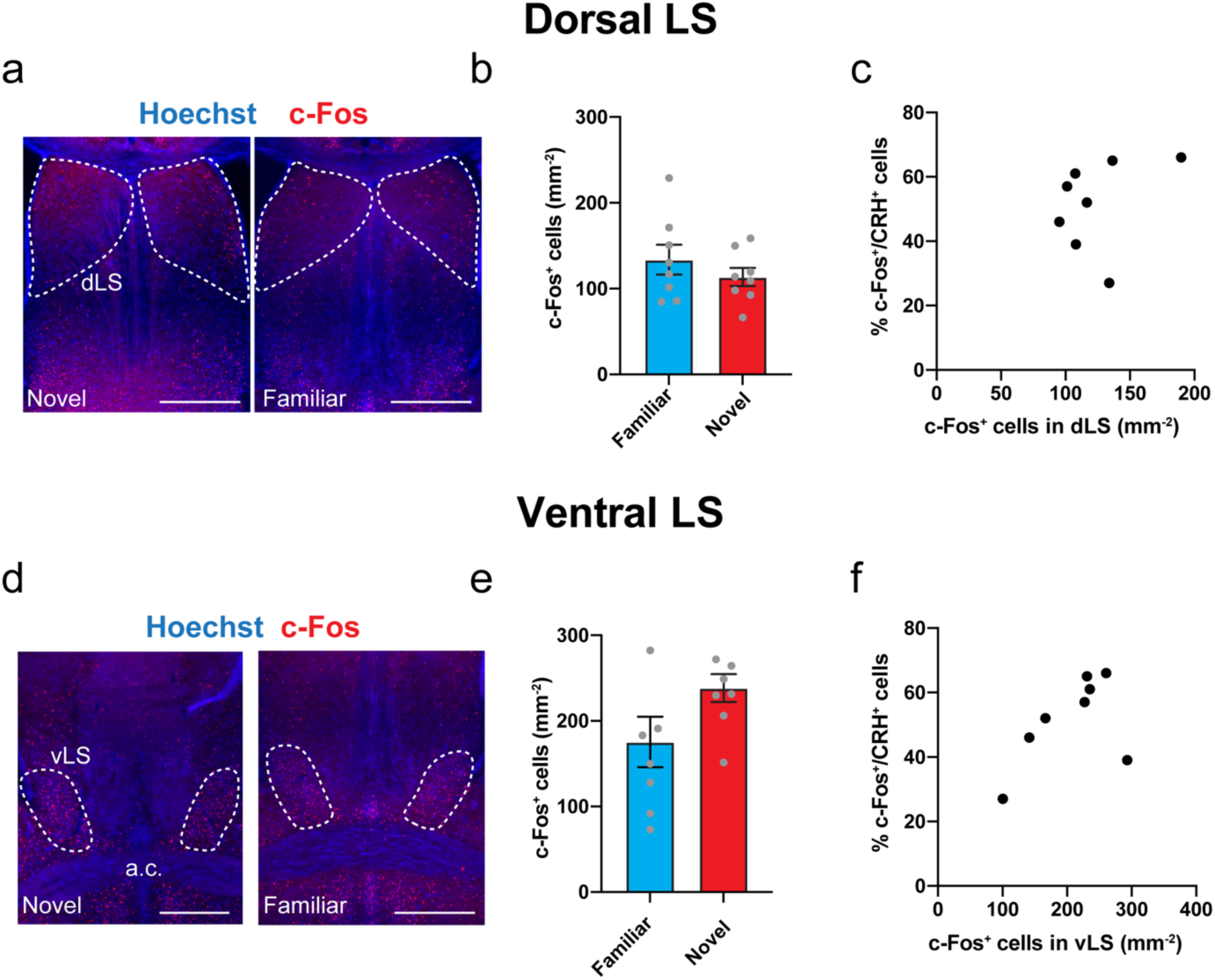
Dorsal and ventral posterior LS do not respond to social familiarity versus social novelty. **a.** Immunohistochemistry images of c-Fos labelling in posterior dorsal LS (dLS). Scale bars: 500 µm. **b.** Density of dLS cells positive for c-Fos. Each point is from a different image obtained from 4 mouse. Unpaired *t* test, *p* = 0.3. **c.** Percentage of ILA^CRH^ cells positive for c-Fos in layer 2/3 vs. density of dLS cells positive for c-Fos following social interaction. Each point represents one mouse. **d.** Immunohistochemistry images of c-Fos labelling in posterior ventral LS (vLS). Scale bars: 500 µm. **e.** Density of vLS cells positive for c-Fos. Each point is from a different image obtained from 4 mouse. Unpaired *t* test, *p* = 0.08. **f.** Percentage of ILA^CRH^ cells positive for c-Fos in layer 2/3 vs. density of vLS cells positive for c-Fos following social interaction. Each point represents one mouse. For the entire figure, bar graphs represent mean ± S.E.M.

**Extended Data Figure 12:**
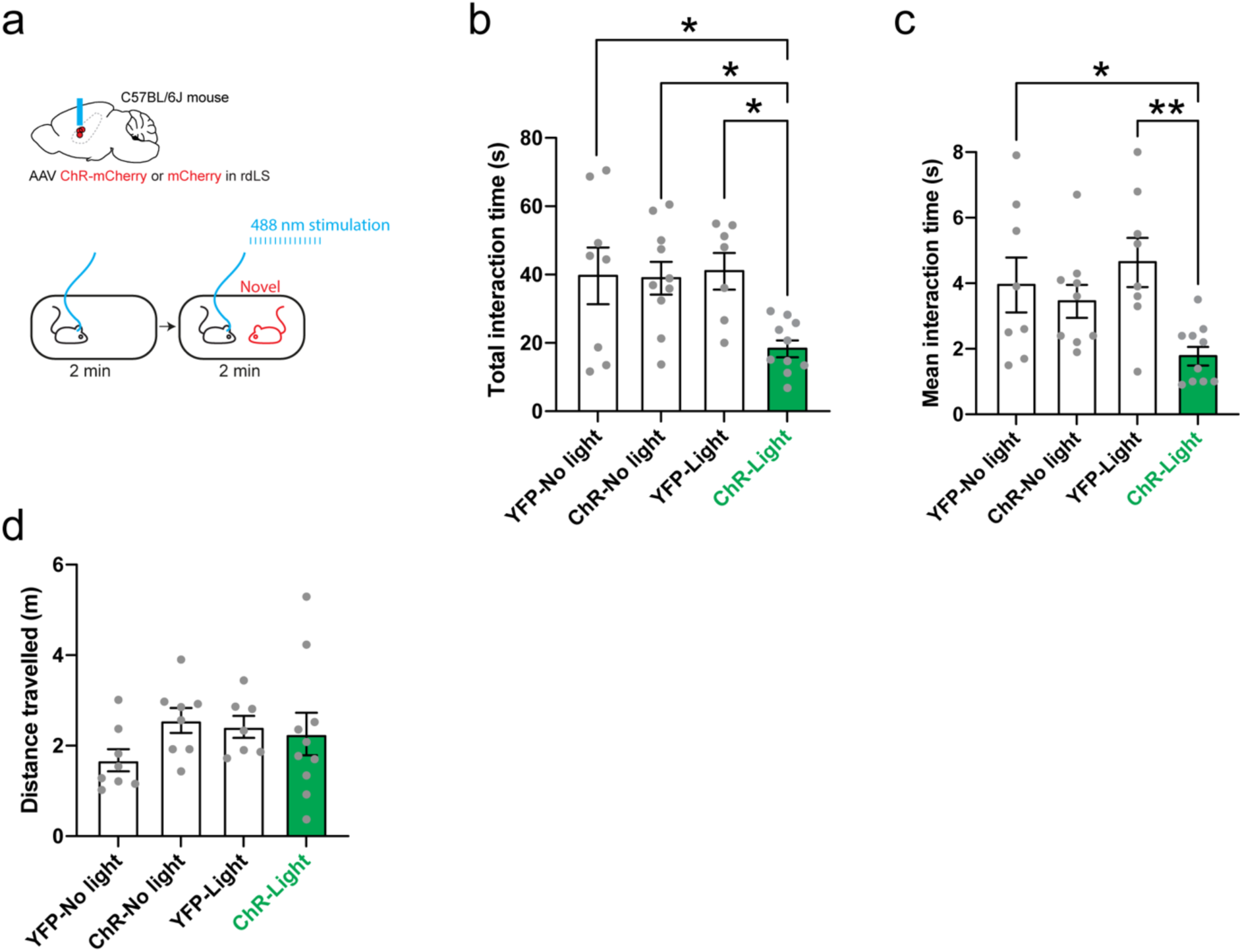
rdLS optogenetic stimulation during interaction with a novel mouse. **a.** C57BL/6J wild-type mice were injected with AA2/2 hSyn1.hChR2(H134R)-mCherry or AA2/2 hSyn1.mCherry as control and an optical fiber was implanted above the injection site. Mice were then presented to a novel mouse for 2 min meanwhile 488 nm light was applied (20 Hz, 1 ms). Mice were also run without light as additional controls. **b.** Total interaction time with novel mouse. Each point represents one mouse. One-way ANOVA: F_3,31_ = 4.47, *p* = 0.01. Dunnett’s multiple comparisons tests: ChR-light vs. YFP-no light *p* = 0.02, ChR-light vs. ChR-no light *p* = 0.02, ChR-light vs. YFP-light *p* = 0.02. **c.** Average duration of each bout of social interaction. Each point represents one mouse. One-way ANOVA: F_3,31_ = 10.62, *p* < 0.0001. Dunnett’s multiple comparisons tests: ChR-light vs. YFP-no light *p* = 0.04, ChR-light vs. ChR-no light *p* = 0.1, ChR-light vs. YFP-light *p* = 0.005. **d.** Total distance travelled. Each point represents one mouse. For the entire figure, bar graphs represent mean ± S.E.M.

**Extended Data Figure 13:**
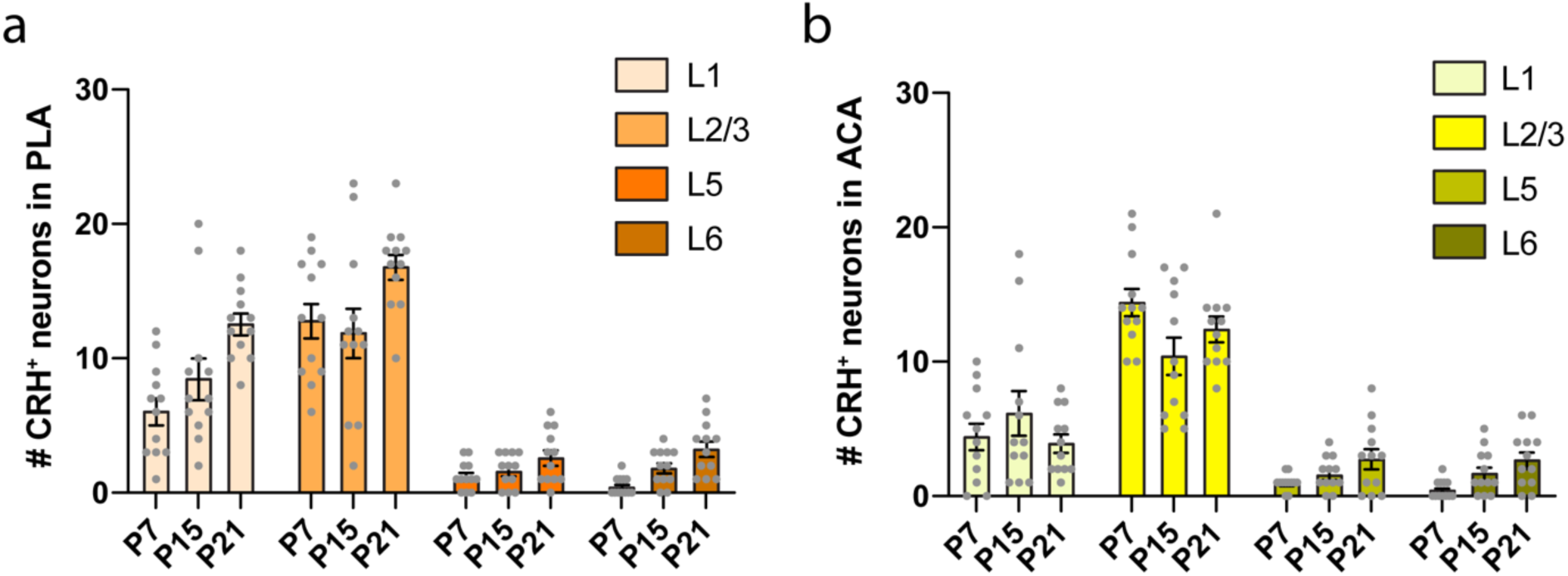
CRH distribution control. **a.** Evolution of the number of CRH^+^ cells per PLA layers. **b.** Evolution of the number of CRH^+^ cells per ACA layers. Each point represents one observation made on each side of 2 section, 3 mice per group. Bar graphs represent mean ± S.E.M.

## Material and methods

### Key resources

| REAGENT or RESOURCE | SOURCE | IDENTIFIER |
| --- | --- | --- |
| **Antibodies** |  |  |
| Anti-GFP antibody produced in chicken | AVES Labs | #GFP-1020 RRID:AB_10000240 |
| Anti-GFP polyclonal antibody, Alexa Fluor 488 | Thermo Fisher Scientific | #A21311<br>RRID:AB_221477 |
| Anti-RFP antibody produced in rabbit | Rockland Antibody | #600-401-379<br>RRID:AB_2209751 |
| Anti-GABA antibody produced in guinea-pig | Abcam | #ab17413<br>RRID:AB_443865 |
| Monoclonal anti-Parvalbumin antibody produced in mouse | Sigma-Aldrich | #P3088<br>RRID:AB_477329 |
| Anti-SST antibody produced in rat | Millipore | #MAB354<br>RRID:AB_2255365 |
| Goat anti-rabbit IgG (H+L) secondary antibody, Alexa Fluor 568 conjugate | Thermo Fisher Scientific | #A11011<br>RRID:AB_143157 |
| Goat anti-rabbit IgG (H+L) secondary antibody, Alexa Fluor 647 conjugate | Thermo Fisher Scientific | #A32733<br>RRID: AB_2633282 |
| Goat anti-guinea-pig IgG (H+L) secondary antibody, Alexa Fluor 647 conjugate | Thermo Fisher Scientific | #A21450<br>RRID:AB_2735091 |
| Goat anti-mouse IgG1 (H+L) secondary antibody, Alexa Fluor 647 conjugate | Thermo Fisher Scientific | #A21240<br>RRID:AB_2536165 |
| Goat anti-rat IgG (H+L) secondary antibody, Alexa Fluor 647 conjugate | Thermo Fisher Scientific | #A48262<br>RRID:AB_2896330 |
| Goat Anti-chicken IgG (H+L) secondary antibody, Alexa Fluor 488 Conjugate | Thermo Fisher Scientific | #A11039<br>RRID:AB_142924 |
| **In situ hybridization probes** |  |  |
| *Mm-Crh* | ACD Bio | #316091 |
| *mCherry-C2* | ACD Bio | #431201-C2 |
| **Chemicals, Peptides, and Recombinant Proteins** |  |  |
| Stressin-1 | Tocris | #1608 |
| Antalarmin | Tocris | #2778 |
| DMSO | Sigma-Aldrich | #D8418 |
| CNO | Cayman Chemical | #16882 |
| **Experimental Models: Organisms/Strains** |  |  |
| C57BL/6J *Mus musculus* | Jackson Laboratories | RRID:IMSR_JAX:000664 |
| B6(Cg)-*Crh*<sup>tm1(cre)Zjh</sup>/J | Jackson Laboratories | RRID:IMSR_JAX:012704 |
| B6-*Crhr1*<sup>tm4(cre)Jde</sup> | Jan Deussing | RRID:MGI:6281608 |
| B6.Cg-*Gt(ROSA)26Sor*<sup>tm9(CAG-tdTomato)Hze/J</sup> | Jackson Laboratories | RRID:IMSR_JAX:007909 |
| **Recombinant DNA** |  |  |
| AAV2/DJ hSyn.FLEX.mGFP.2A.Synatophysin-mRuby | Stanford vector core | #1930 / Addgene #71760 |
| HSV hEF1a.LSIL.GFP (HT) | Massachusetts General Hospital | #RN406 |
| AAV2/8 syn.DIO.TVA.2A.GFP.2A.B19G | University of North Carolina | Addgene #52473 |
| Rabies SAD.B19.EnvA.ΔG.mCherry | Friedrich Miescher Institute for Biomedical Research | Addgene #32636 |
| AAV2/8 hSyn.DIO.hM4D(Gi)-mCherry | Addgene | #44362-AAV8 |
| AAV2/8 hSyn.DIO.mCherry | Addgene | #50459-AAV8 |
| AAV2/5-hSyn-DIO-hM3D(Gq)-mCherry | Addgene | #44361-AAV5 |
| AAV2/5 hsyn.DIO.mCherry | Addgene | #50459-AAV5 |
| AAV2/1 syn.FLEX.GCaMP6f.WPRE.SV40 | Addgene | #100833-AAV1 |
| PX552 | Addgene | #60958 |
| rAAV-U6-shRNA1(CRH)-CMV-EGFP-SV40-polyA | Brain VTA | #PT-2784 |
| rAAV-U6-shRNA2(CRH)-CMV-EGFP-SV40-polyA | Brain VTA | #PT-2785 |
| rAAV-U6-shRNA3(CRH)-CMV-EGFP-SV40-polyA | Brain VTA | #PT-2786 |
| rAAV-U6-shRNA(scramble)-CMV-EGFP-SV40-polyA | Brain VTA | #PT-0916 |
| rAAV-CMV-CRH-P2A-EGFP-WPRE-hGH-polyA | Brain VTA | #PT-2827 |
| AAV2/9 CMV-DIO-(mCherry-U6)-shRNA(anti-*Crh*) | Brain VTA | #PT-2787 |
| AAV2/9 CMV-DIO-(mCherry-U6)-shRNA(scrambled) | Brain VTA | #PT-2788 |
| AAV2/5 DIO.mGFP | University of North Carolina | #AV4310i |
| AAV2/2 CAG.FLEX.ArchT-TdTomato | University of North Carolina | Addgene #28305 |
| AAV2/2 CAG.FLEX.TdTomato | University of North Carolina | Addgene #28306 |
| AAV2/1 syn.GCaMP6f.WPRE.SV40 | Pennsylvania University | #AV-1-PV2822 |
| AA2/2 hSyn1.hChR2(H134R)-mCherry.WPRE | University of Zurich | #V-124 |
| AA2/2 hSyn1.mCherry.WPRE | Addgene | #114472-AAV2 |
| **Software** |  |  |
| AxoGraph | AxoGraph | 1.6.4 |
| PRISM 9 | Graphpad | 9.0.1 (128) |
| Microsoft office Word | Microsoft | 2019 16.56 |
| Microsoft office Exel | Microsoft | 2019 |
| Adobe illustrator Illustrator | Adobe | 2020 v24.1 |
| FIDJI | GPL v2 | 2.3.0/1.53f |
| MATLAB | Mathworks | 2018 |
| Guppy | Lerner Lab <sup>69</sup> | 1.1.4 |
| Leica Application Suite X | Leica |  |
| ANY-maze | Stoelting Co. | 4.99 |

### CONTACT FOR REAGENT AND RESOURCE SHARING

Further information and requests for reagents may be directed to Felix Leroy.

### EXPERIMENTAL MODELS AND SUBJECT DETAILS

All animal procedures were performed in accordance with the regulations of the UMH-CSIC IACUCs. We used P5 to 16-week-old C57BL6/J wild-type (Jackson Laboratories, #000664) mice as well as mice of the same age range from the following transgenic mouse lines: *CRH-Cre* mice (Jackson Laboratories, #012704) and *CRHR1-Cre* mice (courtesy of Jan Deussing). CHR-Cre were crossed to the *Ai9* TdTomato Cre-reporter mice (Jackson Laboratories, #007909) in order to visualize CRH^+^ neurons. All transgenic mice were on the C57BL6/J background. During social interaction tests, stimulus mice were C57BL6/J wild-type mice of the same gender and age than the test mouse. We observed no difference related to sex and the results were pooled together.

## METHOD DETAILS

### Anti-*Crh* shRNA design and in vitro validation

Three different shRNAs that target *Crh* (shRNA1: GCCCTTGAATTTCTTGCAGCC; shRNA2: GCATGGGTGAAGAATACTTCC; shRNA3: GGAAACTGATGGAGATTATCG) were cloned into the PX552 plasmid (Addgene #60958) to make rAAV-U6-shRNA1,2,3(CRH)-CMV-EGFP-SV40-polyA (Brain VTA #PT-2784, #PT-2785 and #PT-2786). In addition, a scrambled shRNA control was cloned into the same plasmid to make rAAV-U6-shRNA(scrambled)-CMV-EGFP-SV40-polyA (Brain VTA #PT-0916). To overexpress CRH mRNA, the sequence for mouse CRH was cloned into a plasmid to construct rAAV-CMV-CRH-P2A-EGFP-WPRE-hGH-polyA (Brain VTA #PT-2827). For validation, HEK293T cells were transfected, in triplicate, with one of the 3 plasmids containing anti-*Crh* shRNAs or the scrambled shRNA construct along with the overexpression plasmid for *Crh* (CMV-CRH-GFP). The cells were collected 48hr post-transfection and RNA was purified (MiniBEST Universal RNA Extraction Kit; Takara, 9767) and subjected to RT-PCR (One Step SYBR®PrimeScript™RT-PCR Kit II; Takara, RR086A) using primers for CRH (F: CCCCGCAGCCCTTGAATTTCTTG; R: GGGCGTGGAGTTGGGGGACAG) and GAPDH (F: GCAAATTCCATGGCACCGTCAAGG; R: CGCCAGCATCGCCCCACTTG) as a control. The RT-PCR results revealed that all three of the anti-*Crh* shRNAs showed robust decreases of *Crh* mRNA relative to the scrambled shRNA control. The anti-*Crh* shRNA-2 showed the highest knockdown efficiency and was selected for in vivo knockdown experiments. The anti-*Crh* shRNA-2 and the scrambled shRNA were cloned into a Cre-dependent plasmid ^70^ to make rAAV-CMV-DIO-(mCherry-U6)-shRNA(*anti-Crh*)-WPRE-hGH-polyA (Brain VTA, #PT-2787) and rAAV-CMV-DIO-(mCherry-U6)-shRNA(scrambled)-WPRE-hGH-polyA (Brain VTA, #PT-2788) which were subsequently packaged into AAV9.

### Virus injections

For all injections, animals were anesthetized using isoflurane and given analgesics. A craniotomy was performed above the target region and a glass pipette was stereotaxically lowered down the desired depth. Injections were performed using a nano-inject II (Drummond Scientific). 23 nL were delivered 10 s apart until the total amount was reached. The pipette was retracted after 5 min. With homozygous animals (C57BL/6J wild-type or *CRH-Cre* mice), injection of the virus injection expressing DREADD, ArchT, shRNA(anti*-Crh*) and their control viruses (fluorophore only) was randomized within each cage.

#### AAVs injections in ILA

We injected AAV2/DJ hSyn.FLEX.mGFP.2A.Synatophysin-mRuby (Addgene #71760 prepared by the Stanford University vector core #1930), AAV2/8 hSyn.DIO.hM4D(Gi)-mCherry (Addgene #44362-AAV8), AAV2/8 hSyn.DIO.mCherry (Addgene #50459-AAV8), AAV2/5-hSyn-DIO-hM3D(Gq)-mCherry (Addgene, #44361-AAV5), AAV2/5 hsyn.DIO.mCherry (Addgene #50459-AAV5), AAV2/1 syn.FLEX.GCaMP6f.WPRE.SV40 (Addgene #100833-AAV1), AAV2/9 CMV-DIO-(mCherry-U6)-shRNA anti*-Crh*), AAV2/9 CMV-DIO-(mCherry-U6)-shRNA(scrambled), AAV2/2 CAG.FLEX.ArchT-TdTomato (Addgene #28305 prepared by the University of North Carolina vector core) and AAV2/2 CAG.FLEX.TdTomato (Addgene #28306 prepared by the University of North Carolina vector core) into the ILA of *CRH-Cre* mice. Injections were done bilaterally with 100 nl injected per site. Injection coordinates were the following (in mm from Bregma): AP: 1.65, ML: ±0.1, DV: -2.6. Viruses expressed for a minimum of 2 weeks.

#### Rabies and AAV helper virus injection in ILA

We delivered 50 nl of a the helper virus AAV2/8 syn.DIO.TVA.2A.GFP.2A.B19G (Addgene #52473 prepared by University of North Carolina vector core) into the ILA of CRH-Cre mouse at the following coordinates AP: 1.65, ML: ±0.1, DV: -2.6 (in mm from Bregma). Following 2 weeks of recovery and AAV expression, a secondary surgery was performed and 300 nl of rabies SAD.B19.EnvA.ΔG.mCherry (Addgene #32636 prepared by the Friedrich Miescher Institute for Biomedical Research vector core) was injected at the same coordinates. Mice were killed 7 d later and the brains cut coronally for imaging.

#### AAV injections in rdLS

We injected hEF1a.LSIL.mCherry, AAV2/5 DIO.mGFP (Massachusetts General Hospital #RN406), AAV2/1 syn.GCaMP6f.WPRE.SV40 (University of Pennsylvania #AV-1-PV2822), AA2/2 hSyn1.hChR2(H134R)-mCherry.WPRE (University of Zurich #V-124) and AA2/2 hSyn1.mCherry.WPRE (Addgene #114472-AAV2) into the rdLS of CRH-Cre, *CRHR1-Cre* or C57BL/6J wild-type mice respectively. Injections were done bilaterally with 100 nl injected per site. Injection coordinates were the following (in mm from Bregma): AP: 0.9, ML: 0.4, DV: -2.8 with a 10° angle. Viruses expressed for a minimum of 2 weeks.

### Optical ferrule implants

Animals were anesthetized using isoflurane and given analgesics. The scalp was removed and we applied Vetbond^TM^ (3M™ #7000002814) along the cut. A craniotomy was performed above the target region and the optical ferrule was lowered until the desired depth. Superglue was applied to hold the lens in position and then dental cement (GC FujiCEM 2) was applied to cover the exposed skull and keep the optical ferrule in position. Animal were allowed to recovered for 5 days before being used. For fiberphotometry imaging of ILA^CRH^ cells in the right hemisphere we implanted the optical ferrule (B280-4419-3, Doric) at the following coordinates: AP: 1.65, ML: 0.1, DV: -2.4.

For fiberphotometry imaging of rdLS cells in the right hemisphere we implanted the optical ferrule at the following coordinates: AP: 0.9, ML: 0, DV: -2.65.

For silencing of ILA^CRH^ cells fibers in rdLS cells we implanted optical ferrules (Thorlab # FT200UMT and # CFLC230-10) bilaterally at the following coordinates: AP: 0.9, ML: ±0, DV: -2.65.

For excitation rdLS cells we implanted the optical ferrule (Thorlab # FT200UMT and # CFLC230-10) at the following coordinates: AP: 0.9, ML: 0, DV: -2.65

### Cannula guide implantation and micro-infusion

The mouse scalp was removed and scored before a hole was drilled (AP +0.9, ML ±0). A cannula guide extending 2.4 mm below the pedestal (Plastics One #C315G 2-G11-SPC) was lowered slowly and kept in place using superglue. The skull was then covered with dental cement (GC FujiCEM 2) and dummy cannulas (Plastics One #C315DC-SPC) were inserted into the guides. The mice were returned to their home cages and left to recover for at least 1 week. For rdLS infusion, mice were placed under light isoflurane anesthesia (2%) and the dummy cannula was removed. A cannula (Plastics One #C315I-SPC) projecting 2.5 mm from the tip of the cannula guide was mounted. 0.6 µL of a solution containing 2 µg of antalarmin dissolved in DMSO or DMSO (Sigma-Aldrich # D8418) only were infused over 5 min using the Fusion 200 syringe pusher (Chemix Inc.) mounted with a 2-µl syringe (Hamilton #88511). The cannula was removed 2 min after the end of the micro-infusion. Mice typically recovered fully from the light anesthesia within 5 min. Mice were returned to their home cages 20 min before the test began.

### Immunohistochemistry (IHC)

Mice were anesthetized using isoflurane then perfused in the heart with 10 mL saline and their brains were quickly extracted and incubated in 4% PFA overnight. After 1 h washing in PBS, 60 µm slices were prepared using a Leica VT1000S vibratome (Leica Biosystems). Unless indicated otherwise, slices were permeabilized for 2h in PBS with 0.5% Triton-X100 (T9284, Sigma-Aldrich) in PBS before being incubated overnight at 4°C with primary antibodies diluted in PBS with 0.5% Triton-X in PBS. The slices were washed in PBS for 1 h then incubated overnight at 4°C with secondary antibodies from Thermo Fisher Scientific at a concentration of 1:500 diluted in PBS with 0.1% Triton-X. Hoechst counterstain was applied (Hoechst 33342 at 1:1000 for 30 min in PBS at RT) prior to mounting the slice using fluoromount (Sigma-Aldrich). Images were acquired using inverted confocal microscopes (lsm 900, Zeiss and SPII, Leica) or an epifluorescent microscope (Thunder, Leica). For post-hoc immunocytochemistry after patch-clamp recordings, slices were fixed for 1 h in PBS with 4% PFA and streptavidin was applied during secondary incubation.

- Fig. 1b-e: For mRuby and GFP labelling, primary incubation was performed overnight at 4°C with rabbit anti-RFP (1:500, Rockland Antibody, #600-401-379) and chicken anti-GFP (1:1000, Aves, #GFP-1020) antibodies. Secondary incubation was performed with anti-rabbit antibody conjugated to Alexa 568 (#A11036) and anti-chicken antibody conjugated to 488 (#A11039).
- Fig. 1n: For GFP labelling, incubation was performed overnight at 4°C with anti-GFP antibody conjugated to Alexa 488 (1:500, Thermo Fisher Scientific #A21311). No immunohistochemistry was performed against mCherry.
- Extended Data Fig. 1e: For GABA labelling, incubation was performed during 2 days at 4°C with an anti-GABA antibody (1:200, Abcam, #ab17413). Secondary incubation was performed with anti-guinea-pig antibody conjugated to Alexa 647 (#A21450). No immunohistochemistry was performed against GFP.
- Extended Data Fig. 1j: For PV labelling, incubation was performed overnight at 4°C with an anti-PV antibody conjugated to Alexa 488 (1:2000, Sigma-Aldrich, #P3088). Secondary incubation was performed with anti-mouse IgG1 antibody conjugated to Alexa 647 (#A21240). No immunohistochemistry was performed against TdTomato.
- Extended Data Fig. 1l: For SST labelling, incubation was performed overnight at 4°C with an anti-SST antibody (1:500, Millipore, #MAB354). Secondary incubation was performed with anti-rat antibody conjugated to Alexa 488 (#A48262). No immunohistochemistry was performed against TdTomato.
- Fig. 2b: For mCherry labelling, primary incubation was performed overnight at 4°C with an anti-RFP antibody (1:500, Rockland Antibody, #600-401-379). Secondary incubation was performed with anti-rabbit antibody conjugated to Alexa 568 (#A11036).
- Fig. 2v, 4o & Extended Data Fig. 5a: For c-Fos labelling, primary incubation was performed overnight at 4°C with an anti-c-Fos antibody (1:1000, Abcam, #ab190289). Secondary incubation was performed with anti-rabbit antibody conjugated to Alexa 647 (#A32733). No immunohistochemistry was performed against mCherry or TdTomato.
- Fig. 3j & 4u: For TdTomato labelling, primary incubation was performed overnight at 4°C with an anti-RFP antibody (1:500, Rockland Antibody, 600-401-379). Secondary incubation was performed with anti-rabbit antibody conjugated to Alexa 568 (#A11036).
- Fig. 3f: For GFP labelling, incubation was performed overnight at 4°C with an anti-GFP antibody conjugated to Alexa 488 (1:500, Thermo Fisher Scientific, #A21311).
- Fig. 4k, 4p & Extended Data Fig. 11a,d: For c-Fos labelling, primary incubation was performed overnight at 4°C with anti-c-Fos antibody (1:1000, Abcam, #ab190289). Secondary incubation was performed with an anti-rabbit antibody conjugated to Alexa 568 (#A11036).

### In situ hybridization (ISH)

Mice were anesthetized using isoflurane then decapitated and their brain quickly extracted. Brains were then immersed in dry-ice cold Butan X for 6 s before being stored at -80°C. 16 µm-thick slices were prepared using a Leica cryostat (CM3050 S, Leica Biosystems) and mounted on *Superfrost Plus* microscope slides (12-550-15, FisherBrand). We processed the slices following the RNAscope Fluorescent Multiplex protocol (ACD Bio) with the probes for *Crh* in C1 (#316091), *mCherry* in C2 (#431201-C2). We applied Protease IV for 1 min and used the Amp4 Alt-A color module. DAPI was apply for 30 s prior to mounting using fluoromount. Images were acquired using an inverted confocal microscope (SPII, Leica).

### In vitro electrophysiological recordings

We prepared coronal brain slices from 8- to 12-week-old C57BL6/J mice. Animals were killed under isoflurane anesthesia by perfusion into the right ventricle of ice-cold solution containing the following (in mM): 125 NaCl, 2.5 KCl, 22.5 glucose, 25 NaHCO_3_, 1.25 NaH_2_PO_4_, 3 Na Pyruvate, 1 Ascorbic acid, 2 CaCl_2_ and 1 MgCl_2_). ACSF was saturated with 95% O_2_ and 5% CO_2_, pH 7.4. Brains were cut into 400 µm slices with a vibratome (VT1200S, Leica) in the same ice-cold dissection solution. Slices were then transferred to an incubation chamber containing the same ACSF solution. The chamber was kept at 34°C for 30 min and then at room temperature for at least 1 h before recording. All experiments were performed at room temperature. Slices were mounted in the recording chamber under a microscope. Recordings were acquired using the Multiclamp 700A amplifier (Molecular Device), data acquisition interface ITC-18 (Instrutech) and the Axograph software. We targeted CA2 PNs based somatic location and size in both deep and superficial layer. Whole-cell recordings were obtained from LS neurons in voltage-clamp mode at -70 mV patch pipette (3–5 MΩ) containing the following (in mM): 135 Cs-gluconate, 5 KCl, 0.2 EGTA-Na, 10 HEPES, 2 NaCl, 5 ATP, 0.4 GTP, 10 phosphocreatine, and 5 µM biocytin, pH 7.2 (280–290 mOsm). The liquid junction potential was 1.2 mV and was left uncorrected. Inhibitory currents were recorded in voltage-clamp mode at +10 mV. After 10 min of stable baseline recording, stressin-1 (300 nM, Tocris # 1608) was applied following a 1:1000 dilution from stock solution into the ACSF. We used the Axograph software for data acquisition, and Excel (Microsoft) and PRISM (Graphpad) for data analysis.

### Behavioral tests

Based on our experience conducting similar social behavior experiments, we used group size of 10-15 animals. Animals that had viral expression outside of ILA or rdLS were excluded from analysis. This criterion was pre-established since we wanted to investigate the role of local neurons. The observer was blind to the identity of the mice while performing the behavioral experiments and the subsequent analyses.

#### Open arena test

Mice were introduced into a new arena (60 cm × 60 cm) and allowed to roam freely for 10 min. Using automatic tracking of the test mouse (Any-Maze 7, Stoelting), we quantified the total distance travelled as well as time spent in the surround (20% of the surface) or center (remaining 80% of the surface).

#### Elevated plus maze test

Mice were placed at the center of a maze consisting of a cross with two open arms and two closed arms. They were allowed to explore the maze freely for 5 min. We quantified the amount of time and number of entries in open or closed arms.

#### Sociability test

Test mice were introduced into a same open arena and allowed to roam freely for 10 min. Then, wire cup cages were introduced at opposite corners. One cup had a novel mouse from the same sex and age underneath it. The test mouse was allowed to explore each cup for 5 min. Using automatic tracking of the test mice (ANY-Maze, Stoelting), we quantified the time spent in the area surrounding each cup. Using the interaction times, we calculated a discrimination index for social preference as the following: DI = (time with mouse – time with empty cup) / total interaction time. DI represents the percentage of extra time spent with the mouse compared to the empty cup. The preference exhibited by the test mouse for the mouse compared to the empty cup was used as an index for sociability.

#### Social novelty preference test

This test was performed in the same arena as the open-arena and sociability test. Test mouse was allowed to explore the arena for 10 min. Then, the test mouse explored for 5 min two wire cups on opposite corners (learning trial). Each cup contained a novel mouse. Stimulus mice were removed from the cups and the test mouse were placed in an empty cage, the size of its home cage for 30 min. Then, one of the now familiar mice was returned under its cup while a novel mouse was introduced under the other cup (recall trial). The test mouse was free to explore each cup again for 5 min. Importantly, all 3 stimulus mice were littermates housed in the same cage (thus preventing the test mice to use any cage-specific odor clue). The design of this test allows for better exploration of each cup than the classical 3-chamber test while still giving the test mouse freedom to explore the novel or familiar mice unlike the direct interaction test. Using automatic tracking of the test mice (ANY-Maze, Stoelting), we quantified the time the test mice spent in the immediate area surrounding each cup - and therefore, interacting with each stimulus mice - during learning and recall. Using the interaction time from the recall trial, we calculated a discrimination index for social novelty preference as the following: DI = (time with novel – time with familiar) / total interaction time. DI represents the percentage of extra time spent with novel compared to familiar. The preference normally exhibited by the test mouse for the novel compared to the familiar animal was used as an index for social memory. We also calculated the total time spent interacting with stimulus mice during each trial in order to verify that the sociability drive was not affected.

#### Object recognition memory test

Same as above but with 3 different small toys of different shape about the size of the mouse. Objects were of the same color.

#### Repetitive social presentation test

Test mice were introduced in a clean cage the same size than their home cage and left to explore for 5 min. Then, a novel mouse of the same age and sex was introduced for four times during 2 min with 10 min inter-trial. For the 5^th^ trial, a novel age- and sex-matched mouse housed in the same cage than the first one was presented. Films were scored offline for social interaction by a trained observer blind to the experimental condition. Sniffing of any part of the body, allo-grooming and close following counted as social interaction. Interaction times were normalized with respect to the first interaction.

#### Repetitive object presentation test

Same as above but with 2 different small toys of different shape about the size of the mouse. Objects were of the same color.

#### Food-seeking test

Mice were food-deprived overnight and then introduce into a large arena (60×60 cm) containing some crushed chow in the middle. We measured the latency to feed and the amount of time spent feeding.

#### One trial presentation of novel or familiar mice

Test mice were individually placed in a clean cage the night before the test. The next day a novel mouse or a littermate was introduced for 1 single trial during 2 min. The total time of interaction during the 2 min was simultaneously annotated during film recording. Social interaction was measure by a key-press using Any-Maze when sniffing of any part of the body or chasing occurred. Mice were perfused 1 hour after performing the test and processed for c-Fos analysis.

#### Kinship preference test

For this test, we used two females with a positive plug from the same day in order to have two litters of the same developmental stage. After the pups of both litters were born, we started to test the percentage of sibling choice of each pup during consecutive days. From P5 to P14 the test was performed as following: each nest containing their corresponding litter was placed in each corner of the OA. Then we place each individual pup in a middle point equidistant from both nests. We recorded each pup during 2 min, then we annotated weather the pup ended reaching the sibling or non-sibling nest. Due to the natural increasing exploration behavior of the pups, after P14, we placed all pups of each litter under a pencil cup and placed them in opposite zones of the OA. Then we tested the percentage of sibling preference by measuring the time spent in each zone during 2 min. We only used litters of 5 pups or more, and with similar number of pups between the litters. We also measured the time spent near each nest to calculate a discrimination index for familiar kin similar to the one calculated previously.

#### Optogenetic stimulation during social interaction

Test mice were introduced in a clean cage the same size than their home cage and left to explore for 5 min. Then, light-stimulation was applied and a littermate or a novel mouse of the same age and sex was introduced for 2 minutes. Films were scored offline for social interaction by a trained observer blind to the experimental condition. Sniffing of any part of the body, allo-grooming and close following counted as social interaction.

### Fiberphotometry data acquisition and analysis

#### Fiber-photometry recording during social or object presentation

Test mice were individually placed in a clean cage the night before the test. The day of recording, mice were habituated to the optical fiber for three days by placing the fiber on the mouse head and letting it roam free for 15 min in their own cage before the test. Then, we recorded basal activity during two minutes. After 5 minutes, we performed 2 min social or object presentations with 5 minutes in between recordings. Novel and familiar mice or objects were introduced for 1 single trial during 2 min. The order of the familiar or novel presentations were randomized along all the sessions. Movies were annotated offline for the beginning and end of the interactions.

#### Fiber-photometry recording during repetitive social presentation

Mice were habituated to the optical fiber for three days by placing the fiber on the mouse head and letting it roam free for 10 min. The test was conducted as described previously.

Fiberphotometry was conducted using a DORIC system (Basic FMC). Two LEDs (405 nm and 465 nm) were coupled to a fluorescence mini-cube (FMC) to deliver light into optical ferrules permanently implanted above the ILA or rdLS. We used 217 Hz as the oscillation frequency for the 465 nm calcium-dependent fluorescence channel, and 319 Hz as the frequency for the 405 nm isosbestic signal. Light was delivered at a final intensity of 2.24 mW (465nm) and 2.76 mW (405 nm) at the tip of the patch-cord before coupling with the implanted ferrule ^71^. Emitted light between 420 and 450 nm (with 405 nm excitation) and 500 and 540 nm (with 465 nm excitation) were collected through the FMC on separate fiber-coupled Newport 2151 photo-receiver modules. The collected fluorescent signals were collected in AC-low mode and converted to voltage via the formula *V* = *PRG*, where *V* is the collected voltage, *P* is the optical input power in watts, *R* is photodetector responsivity in amps/watts (0.2 – 0.4), and *G* is the trans-impedance gain of the amplifier. Raw signals for 473 nm excitation (GCaMP6f) and 405 nm excitation (isosbestic signal) were recorded using Doric Neuroscience Studio software. We used the Guppy software for analysis ^69^. After loading the data, we apply an artifact correction criterion consisting on the removal of the first second of the raw data. This allowed us to remove the artifacts that are usually generated right after the light sources turn on. Subtraction of the background fluorescence was calculated via a time-fitted running average of the 465 nm channel relative to the 405 nm control channel using a least squares polynomial fit of degree 1. Δ*F*/*F* is calculated by subtracting the fitted control channel from the signal channel and dividing by the fitted control channel using the formula (465 nm – 405 nm)/405 nm. A peak enveloping Fourier transform was applied to the Δ*F*/*F* signal across the entire trace to identify peaks in activity. Finally, we presented the data as the deviation of the Δ*F*/*F* from its mean (z-score) using the formula (Δ*F*/*F – mean of* Δ*F*/*F/ Standard deviation of* Δ*F*/*F)*. For PSTH analysis, we analyzed the z-score for each interaction during the sessions based on the following time window (0 to 3 s after the timestamp).

### Classifier analysis

Fiberphotometry recordings from ILA^CRH^ and rdLS were used to discriminate between novel or familiar interactions. On each session, we detected calcium transients and fit support vector machines (SVM) with linear kernel to discriminate whether each transient occurred during a novel or a familiar interaction (Figs. 2S4, 4S2). We used two different approaches to fit the SVMs: individual sessions and pseudo-populations^72,73^. For the individual analysis, we fitted an independent SVM on each individual session (10 sessions for ILA^CRH^ and 10 sessions for rdLS recordings) and evaluated the cross-validated decoding performance per session. Cross-validation was performed by training the classifier on 80% of the dataset and testing the performance on the remaining 20%. Since this partition was random, we repeated this process 20 times and reported the mean. To evaluate the significance of the obtained decoding performance, we built a null-distribution of decoding performances by randomly shuffling the labels for novel or familiar interaction. On each shuffle iteration, we first shuffled the labels and then evaluated the cross-validated performance as in the original dataset (80% train, 20% test, 20 iterations). We repeated this process 1000 times. The reported *p*-value was obtained by comparing the decoding performance of the original dataset with the null distribution. In all cases, the decoding performance was larger than the largest value of the null distribution (*p* < 0.001). For the pseudo-population analysis, for each session we randomly draw (with replacement) 100 trials from familiar and 100 trials from novel interactions. By combining all recording sessions from a particular brain region, we obtained one matrix for training and one matrix for testing (200 trials x 10 sessions) our linear classifiers (SVM). The matrices for training and testing were obtained by sampling with replacement from 80% and 20% of the trials from the original dataset, respectively. Our procedure ensured that there was no overlap between the train and test matrices. Since the procedure to build these two matrices was random, we repeated this process 100 times and reported the mean decoding performance. To evaluate the significance of the obtained decoding performance for the pseudo-population, we also built a null-distribution of decoding performances by randomly shuffling the labels for novel or familiar interaction on each recording session and proceeded as described above (1000 iterations). The reported p-value was obtained by comparing the decoding performance of the original dataset with the null distribution. As for the individual sessions analysis, the decoding performance was larger than the largest value of the null distribution. In Extended Data Fig. 7, the shaded area corresponds to the 2.5 – 97.5 percentile of the null distribution. In all cases, the mean of the null distribution was very close to 50%.

### Statistics

Statistical tests were performed using PRISM (Graphpad). Results presented in the text, figures and figure legends are reported as the mean ± SEM. * is for *p* < 0.05, ** is for *p* < 0.01, *** is for *p* < 0.001 and **** is for *p* < 0.0001.

## Bibliography

1. Amdam, G. V. & Hovland, A. L. Measuring Animal Preferences and Choice Behavior. Nature Education Knowledge 3, 1–9 (2012).

2. Blaustein, A. R. & O’Hara, R. K. Genetic control for sibling recognition? Nature 290, 246–248 (1981).

3. Clemens, A. M., Wang, H. & Brecht, M. The lateral septum mediates kinship behavior in the rat. Nature Communications 11, 3161 (2020).

4. Hepper, P. G. Sibling recognition in the rat. Animal Behaviour 31, 1177–1191 (1983).

5. Kogo, H., Kiyokawa, Y. & Takeuchi, Y. Rats show a preference for unfamiliar strains of rats. bioRxiv 6 (2021) doi:10.1101/2021.02.18.431764.

6. Stowers, L., Holy, T. E., Meister, M., Dulac, C. & Koentges, G. Loss of sex discrimination and male–male aggression in mice deficient for TRP2. Science 295, 1493–1500 (2002).

7. Liu, Y. et al. Molecular regulation of sexual preference revealed by genetic studies of 5-HT in the brains of male mice. Nature 2011 472:7341 472, 95–99 (2011).

8. Kingsbury, L. et al. Cortical Representations of Conspecific Sex Shape Social Behavior. Neuron 107, 941–953.e7 (2020).

9. Mossman, C. A. & Drickamer, L. C. Odor Preferences of Female House Mice (Mus domesticus) in Seminatural Enclosures. Journal of Comparative Psychology 110, 131–138 (1996).

10. Moy, S. S. et al. Sociability and preference for social novelty in five inbred strains: an approach to assess autistic-like behavior in mice. Genes, brain, and behavior 3, 287–302 (2004).

11. Watanabe, S. The dominant/subordinate relationship between mice modifies the approach behavior toward a cage mate experiencing pain. Behavioural Processes 103, 1–4 (2014).

12. Loo, P. L. P. van, Groot, A. C. de, Zutphen, B. F. M. van & Baumans, V. Do Male Mice Prefer or Avoid Each Other’s Company? Influence of Hierarchy, Kinship, and Familiarity. http://dx.doi.org/10.1207/S15327604JAWS0402_1 4, 91–103 (2010).

13. Scheggia, D. et al. Somatostatin interneurons in the prefrontal cortex control affective state discrimination in mice. Nature neuroscience 23, 47–60 (2020).

14. Nadler, J. J. et al. Automated apparatus for quantitation of social approach behaviors in mice. Genes, brain, and behavior 3, 303–314 (2004).

15. Oliva, A., Fernández-Ruiz, A., Leroy, F. & Siegelbaum, S. A. Hippocampal CA2 sharp-wave ripples reactivate and promote social memory. Nature 587, 264–269 (2020).

16. Gheusi, G., Bluthé, R. M., Goodall, G. & Dantzer, R. Social and individual recognition in rodents: Methodological aspects and neurobiological bases. Behavioural Processes 33, 59–87 (1994).

17. Domínguez, S. et al. Maturation of PNN and ErbB4 Signaling in Area CA2 during Adolescence Underlies the Emergence of PV Interneuron Plasticity and Social Memory. Cell Reports 29, 1099–1112.e4 (2019).

18. Laham, B. J., Diethorn, E. J. & Gould, E. Newborn mice form lasting CA2-dependent memories of their mothers. Cell Reports 34, 108668 (2021).

19. Menon, R., Süß, T., Oliveira, V. E. de M., Neumann, I. D. & Bludau, A. Neurobiology of the lateral septum: regulation of social behavior. Trends in Neurosciences vol. 45 27–40 (2022).

20. Bielsky, I. F., Hu, S.-B., Ren, X., Terwilliger, E. F. & Young, L. J. The V1a Vasopressin Receptor Is Necessary and Sufficient for Normal Social Recognition: A Gene Replacement Study. Neuron 47, 503–513 (2005).

21. Lukas, M., Toth, I., Veenema, A. H. & Neumann, I. D. Oxytocin mediates rodent social memory within the lateral septum and the medial amygdala depending on the relevance of the social stimulus: male juvenile versus female adult conspecifics. Psychoneuroendocrinology 38, 916–926 (2013).

22. Gutzeit, V. A. et al. Optogenetic reactivation of prefrontal social neural ensembles mimics social buffering of fear. Neuropsychopharmacology 2020 45:6 45, 1068–1077 (2020).

23. Levy, D. R. et al. Dynamics of social representation in the mouse prefrontal cortex. Nature Neuroscience 2019 22:12 22, 2013–2022 (2019).

24. Kingsbury, L. et al. Correlated Neural Activity and Encoding of Behavior across Brains of Socially Interacting Animals. Cell 178, 429–446.e16 (2019).

25. Park, G. et al. Social isolation impairs the prefrontal-nucleus accumbens circuit subserving social recognition in mice. Cell reports 35, 109104 (2021).

26. Yizhar, O. & Levy, D. R. The social dilemma: prefrontal control of mammalian sociability. Current Opinion in Neurobiology vol. 68 67–75 (2021).

27. Carus-Cadavieco, M. et al. Gamma oscillations organize top-down signalling to hypothalamus and enable food seeking. Nature 542, 232–236 (2017).

28. Vale, W., Spiess, J., Rivier, C. & Rivier, J. Characterization of a 41-residue ovine hypothalamic peptide that stimulates secretion of corticotropin and beta-endorphin. Science (New York, N.Y.) 213, 1394–1397 (1981).

29. Schulkin, J. The CRF signal: Uncovering an information molecule. The CRF Signal: Uncovering an Information Molecule (Oxford University Press, 2017). doi:10.1093/acprof:oso/9780198793694.001.0001.

30. Hostetler, C. M. & Ryabinin, A. E. The crf system and social behavior: A review. Frontiers in Neuroscience 0, 92 (2013).

31. Heinrichs, S. C. Modulation of social learning in rats by brain corticotropin-releasing factor. Brain research 994, 107–114 (2003).

32. Dinan, T. G. et al. Desmopressin normalizes the blunted adrenocorticotropin response to corticotropin-releasing hormone in melancholic depression: evidence of enhanced vasopressinergic responsivity. The Journal of clinical endocrinology and metabolism 84, 2238–2240 (1999).

33. Raadsheer, F. C., Hoogendijk, W. J. G., Stam, F. C., Tilders, F. J. H. & Swaab, D. F. Increased numbers of corticotropin-releasing hormone expressing neurons in the hypothalamic paraventricular nucleus of depressed patients. Neuroendocrinology 60, 436–444 (1994).

34. Risbrough, V. B. & Stein, M. B. Role of Corticotropin Releasing Factor in Anxiety Disorders: A Translational Research Perspective. Hormones and behavior 50, 550 (2006).

35. Dedic, N. et al. Deletion of CRH From GABAergic Forebrain Neurons Promotes Stress Resilience and Dampens Stress-Induced Changes in Neuronal Activity. Frontiers in Neuroscience 13, (2019).

36. Dunn, A. J. & File, S. E. Corticotropin-releasing factor has an anxiogenic action in the social interaction test. Hormones and behavior 21, 193–202 (1987).

37. Elkabir, D. R., Wyatt, M. E., Vellucci, S. V. & Herbert, J. The effects of separate or combined infusions of corticotrophin-releasing factor and vasopressin either intraventricularly or into the amygdala on aggressive and investigative behaviour in the rat. Regulatory peptides 28, 199–214 (1990).

38. Mele, A., Cabib, S., Oliverio, A., Melchiorri, P. & Puglisi-Allegra, S. Effects of corticotropin releasing factor and sauvagine on social behavior of isolated mice. Peptides 8, 935–938 (1987).

39. Gehlert, D. R. et al. Stress and central Urocortin increase anxiety-like behavior in the social interaction test via the CRF1 receptor. European journal of pharmacology 509, 145–153 (2005).

40. Campbell, B. M., Morrison, J. L., Walker, E. L. & Merchant, K. M. Differential regulation of behavioral, genomic, and neuroendocrine responses by CRF infusions in rats. Pharmacology, biochemistry, and behavior 77, 447–455 (2004).

41. Zhao, Y. et al. Subtype-selective corticotropin-releasing factor receptor agonists exert contrasting, but not opposite, effects on anxiety-related behavior in rats. The Journal of pharmacology and experimental therapeutics 323, 846–854 (2007).

42. Bagosi, Z. et al. The effects of CRF and urocortins on the preference for social novelty of mice. Behavioural Brain Research 324, 146–154 (2017).

43. Kasahara, M. et al. Influence of transgenic corticotropin-releasing factor (CRF) over-expression on social recognition memory in mice. Behavioural brain research 218, 357–362 (2011).

44. Chen, P. et al. Prefrontal Cortex Corticotropin-Releasing Factor Neurons Control Behavioral Style Selection under Challenging Situations. Neuron 106, 301–315.e7 (2020).

45. van Pett, K. et al. Distribution of mRNAs encoding CRF receptors in brain and pituitary of rat and mouse. The Journal of comparative neurology 428, 191–212 (2000).

46. Bernardi, S. et al. The Geometry of Abstraction in the Hippocampus and Prefrontal Cortex. Cell 183, 954–967.e21 (2020).

47. Asok, A., Schulkin, J. & Rosen, J. B. Corticotropin releasing factor type-1 receptor antagonism in the dorsolateral bed nucleus of the stria terminalis disrupts contextually conditioned fear, but not unconditioned fear to a predator odor. Psychoneuroendocrinology 70, 17–24 (2016).

48. Wang, Y. et al. Role of Corticotropin-Releasing Factor in Cerebellar Motor Control and Ataxia. Current Biology : CB 27, 2661–2669.e5 (2017).

49. Tanaka, K. Z. et al. Cortical representations are reinstated by the hippocampus during memory retrieval. Neuron 84, 347–354 (2014).

50. Clemens, A. M. & Brecht, M. Neural representations of kinship. Current Opinion in Neurobiology vol. 68 116–123 (2021).

51. Dölen, G., Darvishzadeh, A., Huang, K. W. & Malenka, R. C. Social reward requires coordinated activity of nucleus accumbens oxytocin and serotonin. Nature 501, 179–184 (2013).

52. Hung, L. W. et al. Gating of social reward by oxytocin in the ventral tegmental area. Science (New York, N.Y.) 357, 1406–1411 (2017).

53. Gunaydin, L. A. et al. Natural Neural Projection Dynamics Underlying Social Behavior. Cell 157, 1535–1551 (2014).

54. Li, Y. et al. Serotonin neurons in the dorsal raphe nucleus encode reward signals. Nature communications 7, (2016).

55. Golden, S. A. et al. Basal forebrain projections to the lateral habenula modulate aggression reward. Nature 534, 688–692 (2016).

56. Okuyama, T., Kitamura, T., Roy, D. S., Itohara, S. & Tonegawa, S. Ventral CA1 neurons store social memory. Science 353, 1536–1541 (2016).

57. Phillips, M. L., Robinson, H. A. & Pozzo-Miller, L. Ventral hippocampal projections to the medial prefrontal cortex regulate social memory. eLife 8, (2019).

58. Sun, Q. et al. Ventral Hippocampal-Prefrontal Interaction Affects Social Behavior via Parvalbumin Positive Neurons in the Medial Prefrontal Cortex. iScience 23, (2020).

59. Rawleigh, J. M., Kemble, E. D. & Ostrem, J. Differential effects of prior dominance or subordination experience on conspecific odor preferences in mice. Physiology & behavior 54, 35–39 (1993).

60. Gottfried, J. A. & Zald, D. H. On the scent of human olfactory orbitofrontal cortex: meta-analysis and comparison to non-human primates. Brain research. Brain research reviews 50, 287–304 (2005).

61. Li, W. et al. Right orbitofrontal cortex mediates conscious olfactory perception. Psychological science 21, 1454–1463 (2010).

62. Sun, Q. et al. A whole-brain map of long-range inputs to GABAergic interneurons in the mouse medial prefrontal cortex. Nature neuroscience 22, 1357–1370 (2019).

63. Castelhano-Carlos, M. J., Sousa, N., Ohl, F. & Baumans, V. Identification methods in newborn C57BL/6 mice: A developmental and behavioural evaluation. Laboratory Animals 44, 88–103 (2010).

64. American Psychiatric Association. Cautionary Statement for Forensic Use of DSM-5. in Diagnostic and Statistical Manual of Mental Disorders, 5th Edition (American Psychiatric Publishing, Inc, 2013). doi:10.1176/appi.books.9780890425596.744053.

65. Laryea, G., Arnett, M. G. & Muglia, L. J. Behavioral Studies and Genetic Alterations in Corticotropin-Releasing Hormone (CRH) Neurocircuitry: Insights into Human Psychiatric Disorders. Behavioral sciences (Basel, Switzerland) 2, 135–171 (2012).

66. Aly, M., Yonelinas, A. P., Kishiyama, M. M. & Knight, R. T. Damage to the lateral prefrontal cortex impairs familiarity but not recollection. Behavioural Brain Research 225, 297–304 (2011).

67. Horn, M. et al. The multiple neural networks of familiarity: A meta-analysis of functional imaging studies. *Cognitive*, Affective and Behavioral Neuroscience 16, 176–190 (2016).

68. Moll, J. et al. A neural signature of affiliative emotion in the human septohypothalamic area. The Journal of neuroscience : the official journal of the Society for Neuroscience 32, 12499–12505 (2012).

69. Sherathiya, V. N., Schaid, M. D., Seiler, J. L., Lopez, G. C. & Lerner, T. N. GuPPy, a Python toolbox for the analysis of fiber photometry data. Scientific Reports 2021 11:1 11, 1–9 (2021).

70. Botta, P. et al. Regulating anxiety with extrasynaptic inhibition. Nature Neuroscience 2015 18:10 18, 1493–1500 (2015).

71. Owen, S. F. & Kreitzer, A. C. An open-source control system for in vivo fluorescence measurements from deep-brain structures. Journal of neuroscience methods 311, 170–177 (2019).

72. Boyle, L., Posani, L., Irfan, S., Siegelbaum, S. A. & Fusi, S. The geometry of hippocampal CA2 representations enables abstract coding of social familiarity and identity. bioRxiv 2022.01.24.477361 (2022) doi:10.1101/2022.01.24.477361.

73. Nogueira, R., Rodgers, C. C., Bruno, R. M. & Fusi, S. The geometry of cortical representations of touch in rodents. bioRxiv 2021.02.11.430704 (2021) doi:10.1101/2021.02.11.430704.

